# Insights into opium poppy (*Papaver* spp.) genetic diversity from genotyping-by-sequencing analysis

**DOI:** 10.1101/2021.09.28.462245

**Authors:** Uyen Vu Thuy Hong, Muluneh Tamiru-Oli, Bhavna Hurgobin, Christopher R. Okey, Artur R. Abreu, Mathew G. Lewsey

## Abstract

Opium poppy (*Papaver somniferum*) is one of the world’s oldest medicinal plants and a versatile model system to study secondary metabolism. However, our knowledge of its genetic diversity is limited, restricting utilization of the available germplasm for research and crop improvement. We used genotyping-by-sequencing to investigate the extent of genetic diversity and population structure in a collection of poppy germplasm consisting of 91 accessions originating in 30 countries of Europe, North Africa, America, and Asia. We identified five genetically distinct subpopulations using discriminate analysis of principal components and STRUCTURE analysis. Most accessions obtained from the same country were grouped together within subpopulations, likely a consequence of the restriction on movement of poppy germplasm. Alkaloid profiles of accessions were highly diverse, with morphine being dominant. Phylogenetic analysis identified genetic groups that were largely consistent with the subpopulations detected and that could be differentiated broadly based on traits such as number of branches and seed weight. These accessions and the associated genotypic data are valuable resources for further genetic diversity analysis, which could include definition of poppy core sets to facilitate genebank management and use of the diversity for genetic improvement of this valuable crop.

## Introduction

Opium poppy (*Papaver somniferum* L.) is one of the oldest cultivated plant species. Archaeological evidence shows that poppy has been cultivated and used for thousands of years, dating back to the earliest Neolithic ages^1–4^. However, its origin and domestication history has remained unclear until recently. Domestication traits are poorly defined, making it difficult to distinguish domesticated and wild forms especially in archaeological records^5^. Several lines of evidence, based mainly on archaeological data and geographical distribution of cultivated and wild species, suggest the Mediterranean as the centre of poppy origin and domestication^5,6^. Changes in capsule and seed sizes and capsule indehiscence, which is the retention of seed in the capsules, are believed to be amongst poppy domestication-related traits^7^.

Currently, poppy is widely cultivated as both a licit and illicit crop in Asia, Europe, Oceania and South America^8–10^. It is a source of several benzylisoquinoline alkaloids (BIAs) including morphine, codeine, thebaine, papaverine and noscapine for the pharmaceutical industry and for the clandestine production of heroin. Poppy seeds are also used in the food industry for baking and extraction of edible oil, whilst the plant is grown for ornamental purposes in some countries due to its attractive flowers^11^. Poppy cultivars used in food applications are required to contain no or negligible amounts of alkaloids^12^. The availability of commercial cultivars with specific alkaloid profiles is also vital to meet the needs of the pharmaceutical industry and subsequent consumers^12^. Obtaining codeine and thebaine from morphine-free plants can also contribute to preventing illicit production of the morphine-derived heroin.

Considerable poppy genetic diversity has been reported in several countries including India, Turkey, Czech Republic, and Australia^13–17^. Germplasm collections of varying sizes exist in some of these countries^14,15,18,19^. Additionally, a substantial number of poppy genetic resources are currently maintained as seeds in global genebanks. The Leibniz Institute of Plant Genetics and Crop Plant Research (IPK) genebank in Germany has over 1,100 accessions of poppy that were collected worldwide^20^. A collection of similar size is maintained at the Institute of Protection of Biodiversity and Biological Safety in the Slovak University of Agriculture^21^. Germplasm collections provide the genetic and phenotypic diversity used in crop breeding and development. They are also vital resources for research aimed at dissecting the genetic and molecular basis of essential plant processes including secondary/specialized metabolism. However, these resources are underexploited in poppy partly because the genetic diversity of the available germplasm has not been studied in detail. Additionally, the legal opium poppy industry is strictly regulated under international law, restricting the movement of germplasm between countries. Most available reports of opium poppy genetic diversity are based on studies that either assessed small collections of germplasm from a single country or used a limited number of classical DNA markers such as amplified fragment length polymorphisms (AFLPs), random amplified polymorphic DNAs (RAPDs) and simple sequence repeat markers (SSRs)^13,14,22,23^.

Molecular markers have been instrumental for the study of genetic diversity and population structure of germplasm collections. Such studies generate information key to both germplasm conservation and use of the resources for genetic improvement of crops. As the most abundant types of sequence variation in plant genomes, single nucleotide polymorphisms (SNPs) are suitable for several applications that require high-density and genome-wide markers including genetic diversity and population structure analyses, QTL mapping and map-based cloning^24,25^. Recent advances in next generation sequencing have greatly reduced the cost of genome sequencing, allowing the generation of very large numbers of molecular markers. However, reduced representation sequencing (RRS) is still widely used when analyzing large number of samples in species with large genomes, such as opium poppy. Genotyping-by-sequencing (GBS) is a low-cost and fast RRS method for SNP discovery and mapping, which reduces the complexity of genomes by generating smaller fragments *via* restriction digestion^26^. A draft of the *P. somniferum* genome that captured 94.8% of the estimated genome size, with 81.6% of the sequences assigned to individual chromosomes, has been sequenced recently^27,28^. Given the genome is an estimated 2.8 Gb and comprised of 70% repetitive elements, RRS methods are an appropriate, cost-effective method for poppy diversity analyses. Such studies could facilitate the development of poppy core sets, enabling the mapping and isolation of genes or genomic regions associated with traits of interest.

In this work, we applied GBS analysis to characterize 91 poppy accessions obtained from the IPK genebank in Germany. The accessions consisted of two *Papaver* species, *P. somniferum* L. subsp. *somniferum* (*P*. *somniferum* hereafter) and *P. somniferum* L. subsp. *setigerum* (DC.) Arcang. (*P. setigerum* hereafter). *P. setigerum* is commonly thought to be the direct ancestor of *P. somniferum* and historically has also been used for alkaloid production^29^. The accessions originated from diverse geographic regions that encompassed 30 countries of Europe, North Africa, America, and Asia. We provide a genome-wide assessment of the genetic diversity and population structure of the accessions using GBS. The data generated here provides a resource that will allow detailed analysis of poppy germplasm and the definition of poppy core set for effective management of poppy genetic resources. It may also facilitate the genetic improvement of opium poppy through the generation of mapping populations, the identification of useful traits and development molecular markers for marker-assisted selection.

## Results

### Optimization of a poppy GBS protocol for SNP discovery

To generate a SNP dataset sufficient for genetic diversity analysis, we first optimized a GBS protocol for opium poppy. GBS remains a method of choice for genome-wide SNP detection in non-model species and species with large genomes. It is however yet to be utilized for opium poppy. A critical step in GBS protocols is the reduction of genome complexity using restriction enzymes (REs), and the two-enzyme GBS protocol uses a combination of a rare- and common-cutting REs^30^. Although some RE combinations are often used in plants, the optimal combinations need to be determined for the genome of each species^31^. To select the optimal enzyme combination for opium poppy, we prepared eight double digested libraries from a pool of 3 representative samples using the enzyme combinations *Pst*I/*Msp*I, *Pst*I/*Mse*I, *Pst*I/*Nla*III, *Pst*I/*Hpy*CH4IV, *Eco*RI/*Msp*I, *Eco*RI/*Mse*I, *Eco*RI/*Nla*III and *Eco*RI/*Hpy*CH4IV. We then compared the absence of visible repeat regions within the size selection area (280 bp-375bp) and level of amplifications to select *Eco*RI/*Nla*III as the optimal combination for opium poppy GBS library preparation (Fig. **S1**).

The multiplexed pool of *Eco*RI/*Nla*III-based GBS libraries for 91 accessions was sequenced to generate a total of 103,802,122 raw 150-bp single end reads (15.57 Gb data, average 1.14 million reads per sample, Table **S1,2**). We aligned the 103,601,279 reads to the poppy reference genome after filtering, with a read alignment rate of 97% to 99% (Table **S3**). A total of 165,363 SNPs was identified at 76,407 loci and present in ≥90% of accessions, which is significantly higher than the number of SNPs we were able to call using a reference-free pipeline (Table **S4**) These SNPs were evenly distributed across the 11 chromosomes and unplaced scaffolds of the draft opium poppy reference genome (Fig. **1a,b**). The 165,363 SNPs were predicted to have 165,786 effects, of which 149,536 (90.2%) were found in intergenic regions, while the remaining 16,250 (9.8%) corresponded to genic regions (Table **S5**). This optimized protocol will allow researchers to rapidly apply GBS to unlimited number of poppy accessions at reduced cost, allowing detailed characterization of the available germplasm.

**Fig. 1.**
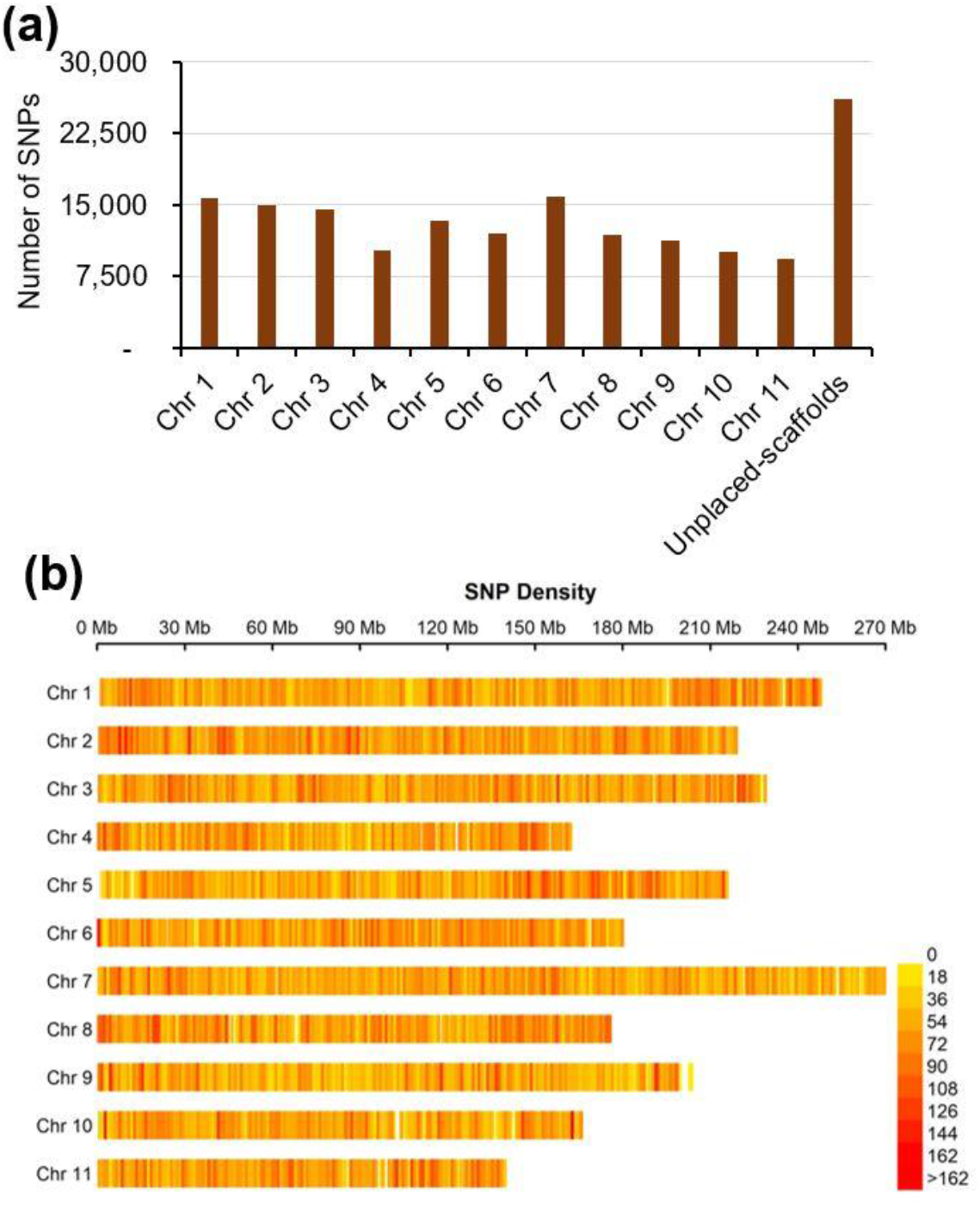
Distribution and density of poppy (*Papaver*) GBS SNPs across the poppy genome. **(a)** Distribution of the of 165,363 SNP markers identified across the 11 chromosomes and unplaced scaffolds of *P. somniferum*. **(b)** SNP density across the 11 *P. somniferum* chromosomes. Number of SNPs in each 1 Mb non-overlapping window are shown.

### Assessing genetic relatedness of 91 *Papaver* accessions

Next, we set out to determine the relationship amongst the 91 *Papaver* accessions from a broad geographic range (Table **S1**). The accessions, which originated from 30 countries in four continents, were primarily *P. somniferum* (88 accessions) but included 3 *P. setigerum* accessions (Table **S1**). We first used pairwise comparisons of the 165,363 filtered SNPs for hierarchical clustering based on the identity-by-state algorithm^32^. Four clusters plus a single accession (PAP 400) far from all others were identified (Fig. **2**). Cluster 1 contained the three *P. setigerum* accessions, while the *P. somniferum* accessions, except PAP 400, were grouped into three distinct clusters. Although PAP 400 was labelled as a *P. somniferum* accession, our result suggests that it is neither *P. somniferum* nor *P. setigerum*. We assessed the morphological characteristics of the accessions when grown under controlled glasshouse conditions. We found that a range of morphological traits were quite diverse between accessions, including seed and capsule characteristics (Fig. **3**; Table **S6;** Fig. **S2-S4**). PAP 400 was morphologically distinct from both *Papaver* species (Fig. **4a**).

**Fig. 2.**
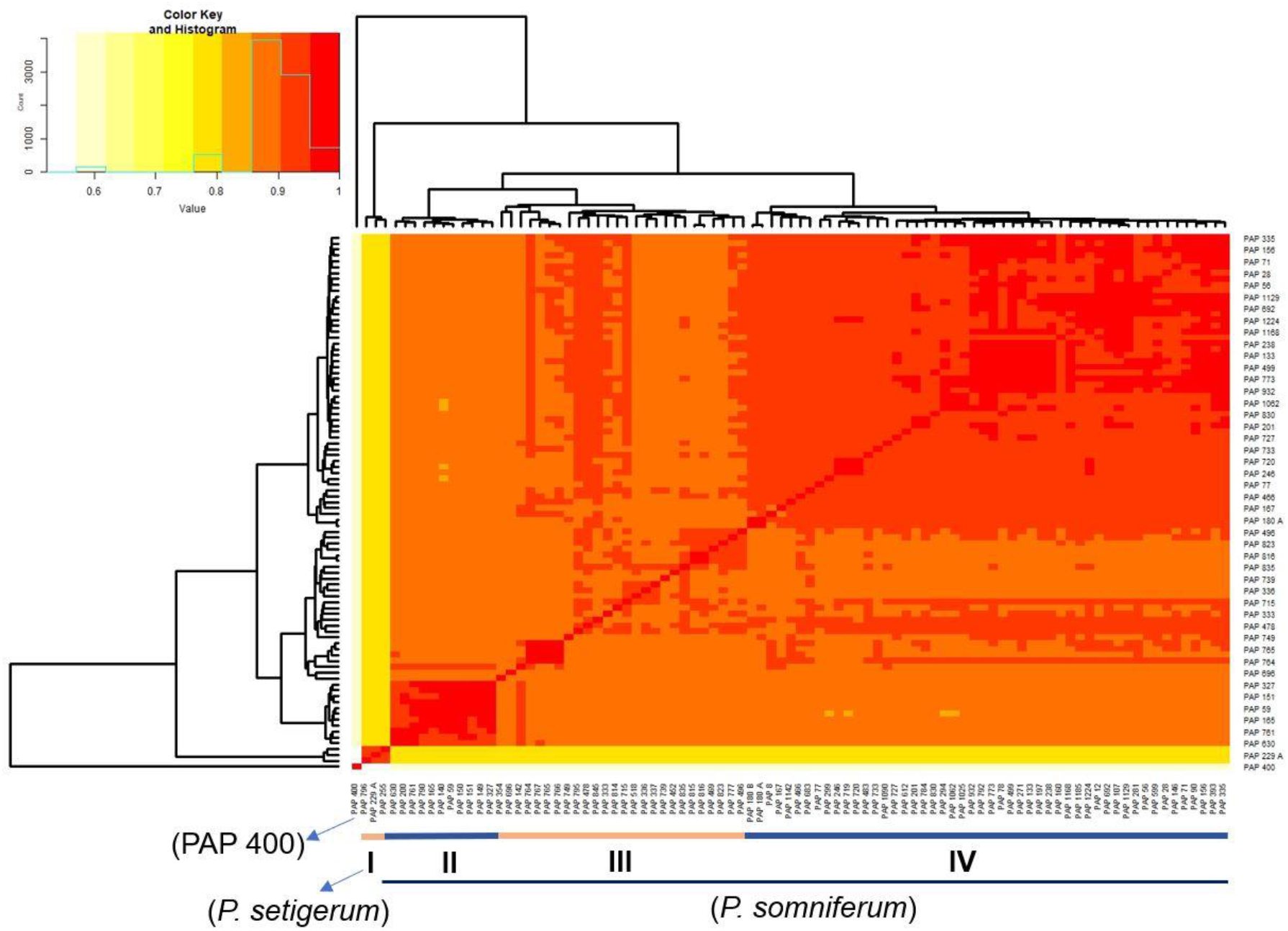
Hierarchical cluster dendrogram and heatmap based on pairwise identity-by-state (IBS) values from 165,363 SNP data representing the genetic relationships among 91 *Papaver* accessions. The four clusters identified are shown with numbers from I - IV. Degree of relatedness is indicated by colours from light yellow (no relatedness) to red (strong relatedness).

**Fig. 3.**
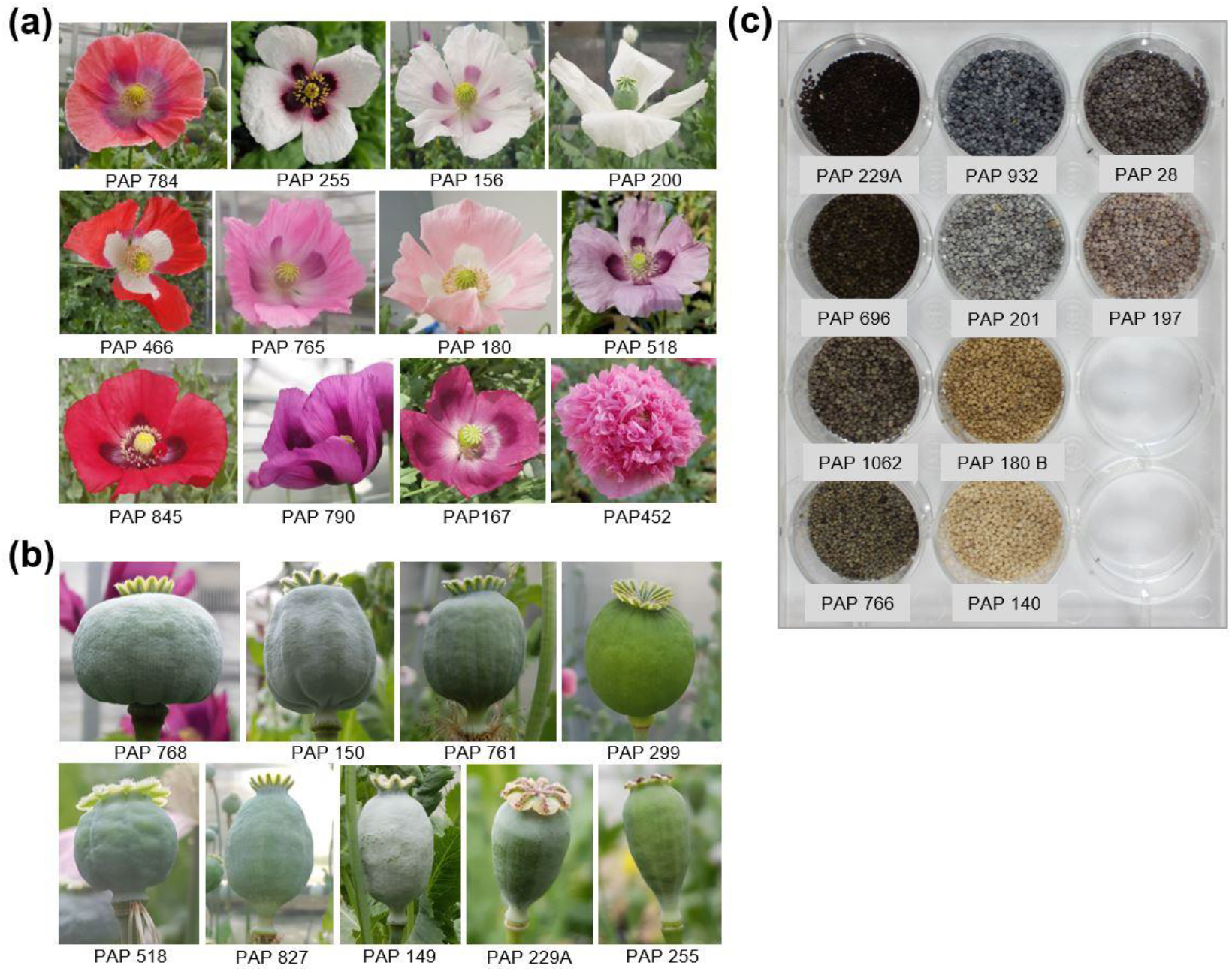
Morphological diversity among the studied poppy (*Papaver*) accessions. Representative images showing **(a)** the distinct floral morphologies, **(b)** the diversity in capsule shape and size and (c) the different seed colour types recorded for the accessions. Accession names are indicated.

**Fig. 4.**
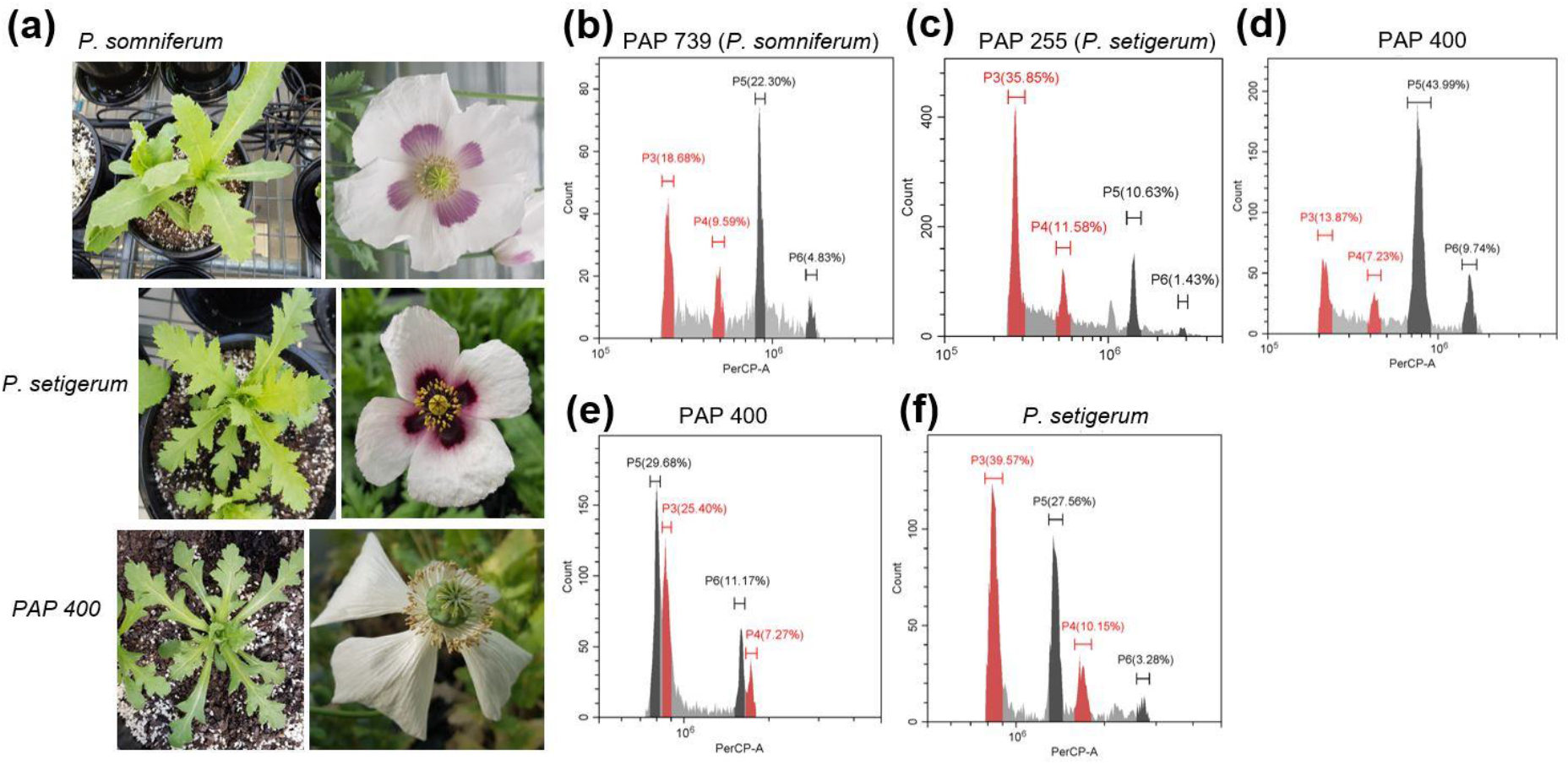
PAP 400 is morphologically distinct from *P. somniferum* and has a smaller genome. **(a)** Leaf and flower morphology of PAP 400, *P. somniferum* and *P. setigerum.* **(b-d)** Fluorescence histograms of PI-stained nuclei from *Papaver* and tomato (internal reference standard, genome size of ~900 Mb) leaves isolated, stained and analysed simultaneously. G1 and G2 peaks of PAP 400, PAP 739 (*P. somniferum*) and PAP 255 (*P. setigerum*) are shown in black, while G1 and G2 peaks of tomato are shown in red. **(e,f)** Fluorescence histograms of PI-stained nuclei isolated from PAP 400 and *P. setigerum* with PAP 518 (*P. somniferum*) used as internal standard. G1 and G2 peaks of PAP 400 and *P. setigerum* were shown in black. The red histograms represent G1 and G2 peaks of *P. somniferum* (internal standard).

To study the relationships between *P. somniferum*, *P. setigerum* and PAP 400, we determined genome sizes and ploidy levels of PAP 400 and representative *P. somniferum* and *P. setigerum* accessions by flow cytometry. We estimated the genome size of *P. somniferum* to be ~3.04 Gb, which is slightly bigger than a previous ~2.87 Gb estimate^27^ (Fig. **4b**). The genome of *P. setigerum* was estimated to be ~4.9 Gb, indicating the *P. setigerum* genome is close to twice the genome size of *P. somniferum* (Fig. **4c,f**). This was similar to previous genome size estimates and supports reports that *P. setigerum* is tetraploid (2n = 4× = 44) with chromosomes smaller in size compared with the diploid (2n = 2× = 22) *P. somniferum*^33–35^. PAP 400 had a slightly smaller genome size than the diploid *P. somniferum* (Fig. **4d,e**). Taken together, our results suggest PAP 400 may be a different species and a case of mislabelling, which can occur in seedbanks during plant cultivation and storage^36^. This is possible given that IPK holds seeds of other *Papaver* species in its collection. However, the dataset we present is relatively small, including only three *P. setigerum* accessions. Consequently, accurate classification of PAP 400 and the accessions in general would require a larger dataset, in particular covering more *P. setigerum* accessions and the other known *Papaver* species.

### Population structure and genetic diversity amongst *Papaver* accessions

Understanding the genetic structure of populations is useful for germplasm conservation and plant breeding. To infer population structure, we analysed the 90 *Papaver* accessions, removing the outlier PAP 400 and associated data (131,039 SNPs remaining). Five clusters (hereafter termed subpopulations) were inferred at the lowest Bayesian information criterion (BIC) score (Fig. **5a**). To understand the genetic relationships between the five subpopulations, we carried out DAPC. Ten principal components (PCs) were retained (with 47.75% of the variance conserved) by the cross-validation function, which gave four discriminant eigenvalues (Fig. **5b**). Subpopulation 1 (SP1) was comprised of all three *P. setigerum* accessions, while SPs 2-5 consisted of 12, 4, 21, and 50 *P. somniferum* accessions, respectively. The wide separation between SP1 and the other subpopulations (SPs 2-5) on the DAPC plot illustrates the extensive genetic difference between *P. somniferum* and *P. setigerum*.

**Fig. 5.**
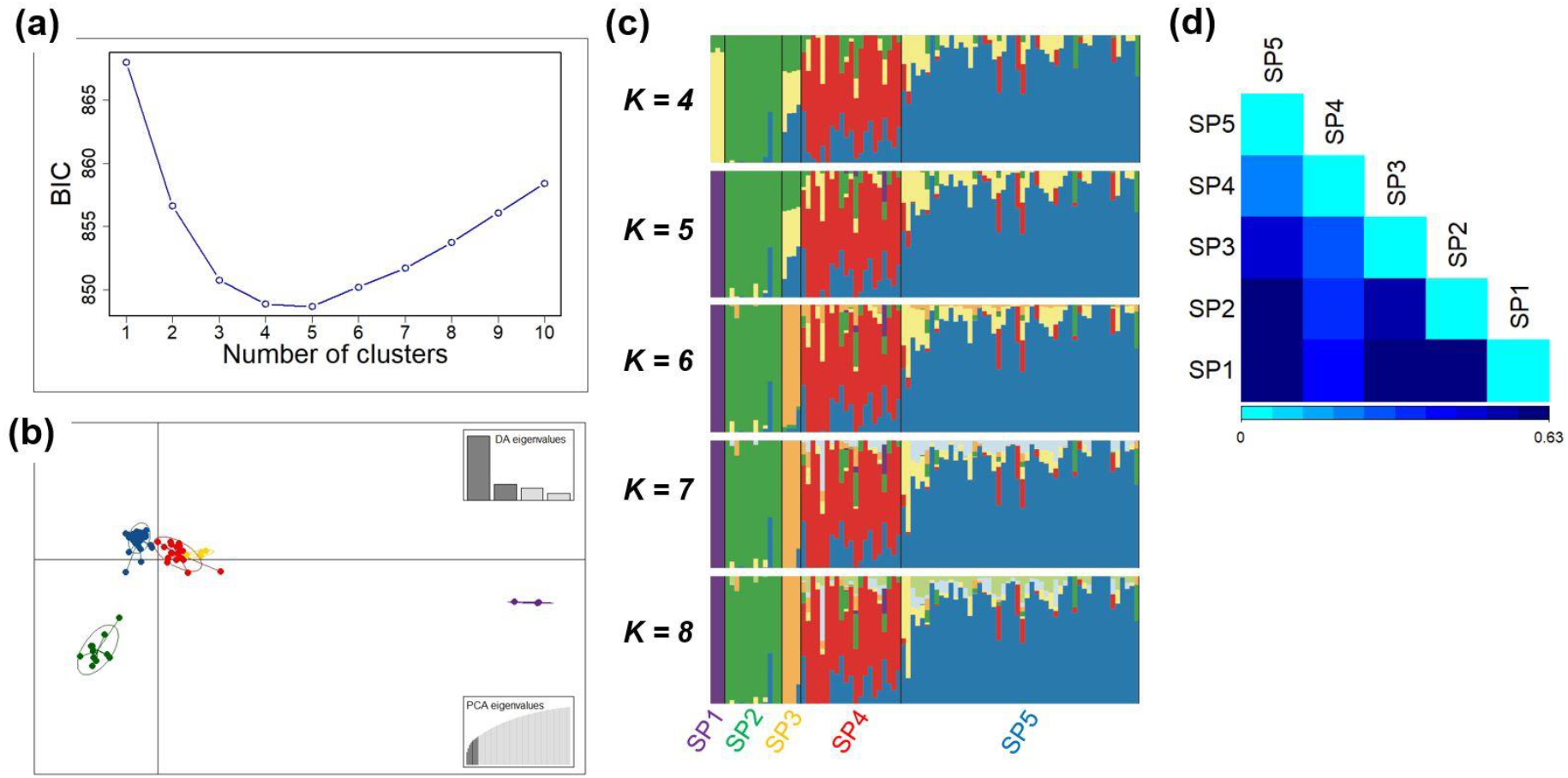
Population structure and differentiation of 90 *Papaver* accessions. **(a)** Bayesian Information Criterion (BIC) values for different number of clusters (subpopulations) using a dataset of 131,039 SNPs. **(b)** Discriminant analysis of principal components (DAPC) for 90 *Papaver* accessions using a dataset of 131,039 SNPs. Ten PCs and four discriminant eigenvalues were retained during analyses to describe the relationship between the subpopulations. The axes represent the first two Linear Discriminants (LD). Each circle represents a subpopulation, and each dot represents an accession. The different colours represent the five subpopulations. **(c)** Genetic structure of 90 *Papaver* accessions estimated by STRUCTURE using various K values. Each accession is represented by a vertical bar divided into K-coloured segments that represent the accession’s estimated common fraction in the K clusters. The accessions are sorted according to DAPC clusters. **(d)** Pairwise population differentiation estimated using *F*_ST_. SP, subpopulation.

We investigated population structure in greater detail. We detected admixture amongst the 90 poppy accessions by applying the admixture model in STRUCTURE using 49,166 unlinked SNPs^37^. Based on ΔK values, the most optimum K value detected was four to eight (Fig. **S5**). Bar plots for each optimal K value, with the accessions sorted following the DAPC result, illustrated that the pattern of subpopulation assignment did not change significantly across the different K values and was consistent with the five subpopulations determined by DAPC (Fig. **5c**). At K = 5, the three *P. setigerum* accessions making up the SP1 from DAPC are genetically distinct with no admixture from the other subpopulations (Fig. **5c**). Interestingly, the 12 accessions of SP2 were a less genetically diverse group representing broad geographic origins extending from North Africa to East Asia, indicative of germplasm exchanges in the past. The four accessions of SP3, which were all from North Korea, had moderate level of admixtures from SPs 2 and 5. The 21 accessions making up SP4 were highly diverse with a high level of admixtures from SPs 2, 3 and 5. Most of the accessions in SP4 originated from western and Mediterranean regions of Europe. Considering that opium poppy was domesticated in the western Mediterranean, from where it spread to north and central Europe, this admixture might be due to ongoing gene flow between wild and domesticated forms. SP5 contained 50 accessions with low level of admixtures from all the other subpopulations^5,6^. Subpopulations were significantly differentiated, as shown by pairwise calculation of genetic differentiation or fixation index (*F*_ST_) that ranged from 0.235 to 0.627, suggesting either low levels of allele sharing or differences in allele frequencies between subpopulations (Fig. **5d**; Table **1**). SP1, comprised of *P. setigerum* accessions, was strongly differentiated from sub-populations of *P. somniferum* accessions, supporting the results of DAPC and STRUCTURE analysis (Fig. **5b,c**).

**Table 1.** Pairwise genetic differentiation (*F*_ST_) between five subpopulations of opium poppy accessions calculated from 49,166 single nucleotide polymorphism loci. *F*_ST_ values are given below the diagonal and those in bold represent significant differences. Corrected *P* values are given above the diagonal (*, α ≤ 0.005, with Bonferroni correction for multiple comparisons).

| Subpopulation | SP1 | SP2 | SP3 | SP4 | SP5 |
| --- | --- | --- | --- | --- | --- |
| SP1 |  | 0.005 | 0.033 | 0.002 | 0.001 |
| SP2 | 0.587 |  | 0.001 | 0.001 | 0.001 |
| SP3 | 0.591 | 0.550 |  | 0.001 | 0.001 |
| SP4 | 0.407 | 0.371 | 0.272 |  | 0.001 |
| SP5 | 0.627 | 0.571 | 0.447 | 0.235 |  |

SP, subpopulation

There were noticeable differences in genetic diversity between the five DAPC subpopulations (Table **2**). The number of private/unique alleles (AP) ranged from 790 (SP3) to 37,157 (SP1; *P. setigerum*), calculated from the 131,039 SNP dataset. This further confirmed the genetic distinctiveness of *P. setigerum*. The percentage of polymorphic loci varied from 3.93% (SP3) to 25.05% (SP5). The level of observed heterozygosity (*H_O_*) was highest for SP1 (0.166) and lowest for SP2 (0.009). *H_O_* was lower than the expected heterozygosity (*H_E_*) for all subpopulations except SP1 (*P. setigerum*), indicating high levels of inbreeding in *P. somniferum*. This finding was supported by the higher inbreeding coefficients (*F_IS_*) for the *P. somniferum* subpopulations (0.061 to 0.377). *F_IS_* was negative for SP1, possibly a consequence of an excess of the observed heterozygotes. The highest nucleotide diversity (*π*) was observed in SP1 (0.219) and the lowest in SP3 (0.040). The genetic variations were both due to differences between (41.6%) and within (44.8%) subpopulations, determined using analysis of molecular variance (AMOVA; Table **3**).

**Table 2.** Measures of diversity for 90 *Papaver* accessions from five subpoulations calculated from 49,166 single nucleotide polymorphism loci.

| Subpopulation | N | AP | %Poly | $H_O$ | $H_E$ | $\pi$ | $F_{IS}$ |
| --- | --- | --- | --- | --- | --- | --- | --- |
| SP1 | 3 | 37,157 | 8.15 | 0.231 | 0.166 | 0.219 | -0.015 |
| SP2 | 12 | 3,937 | 11.54 | 0.009 | 0.061 | 0.064 | 0.189 |
| SP3 | 4 | 790 | 3.93 | 0.009 | 0.034 | 0.040 | 0.061 |
| SP4 | 21 | 14,335 | 24.28 | 0.018 | 0.126 | 0.130 | 0.377 |
| SP5 | 50 | 15,714 | 25.05 | 0.016 | 0.076 | 0.077 | 0.313 |

SP, subpopulation; N, number of individuals; AP, private alleles; %Poly, percentage of polymorphic loci; $H_O$ , observed heterozygosity; $H_E$ , expected heterozygosity; $\pi$ , nucleotide diversity; $F_{IS}$ , inbreeding coefficient

**Table 3.** Analysis of molecular variance for 90 opium poppy accessions based on 49,166 single nucleotide polymorphism markers.

| Source of variation | % Variation | $F$-value | $P$-value |
| --- | --- | --- | --- |
| Between subpopulations | 41.6 | 0.416 | 0.001* |
| Between accessions within subpopulations | 44.8 | 0.767 | 0.001* |
| Within accessions | 13.6 | 0.864 |  |

\* $\alpha \leq 0.05$

We further investigated patterns of accession groupings with regard to their country of origin by combining the hierarchical clustering and population structure analyses with information on the country of origin of each accession (Fig. **6**). The North Korean accessions formed a distinct subpopulation within this analysis. Additionally, some accessions obtained from the same countries were grouped together as genetically similar. Examples included accessions from Japan, Morocco, Belgium, Switzerland, Germany, Mongolia, Russia, Australia, Bulgaria, and Czechoslovakia (Fig. **6**). We interpret these results as reflecting the highly restricted movement of poppy germplasm between different countries and the controlled circumstances under which poppies are cultivated.

**Fig. 6.**
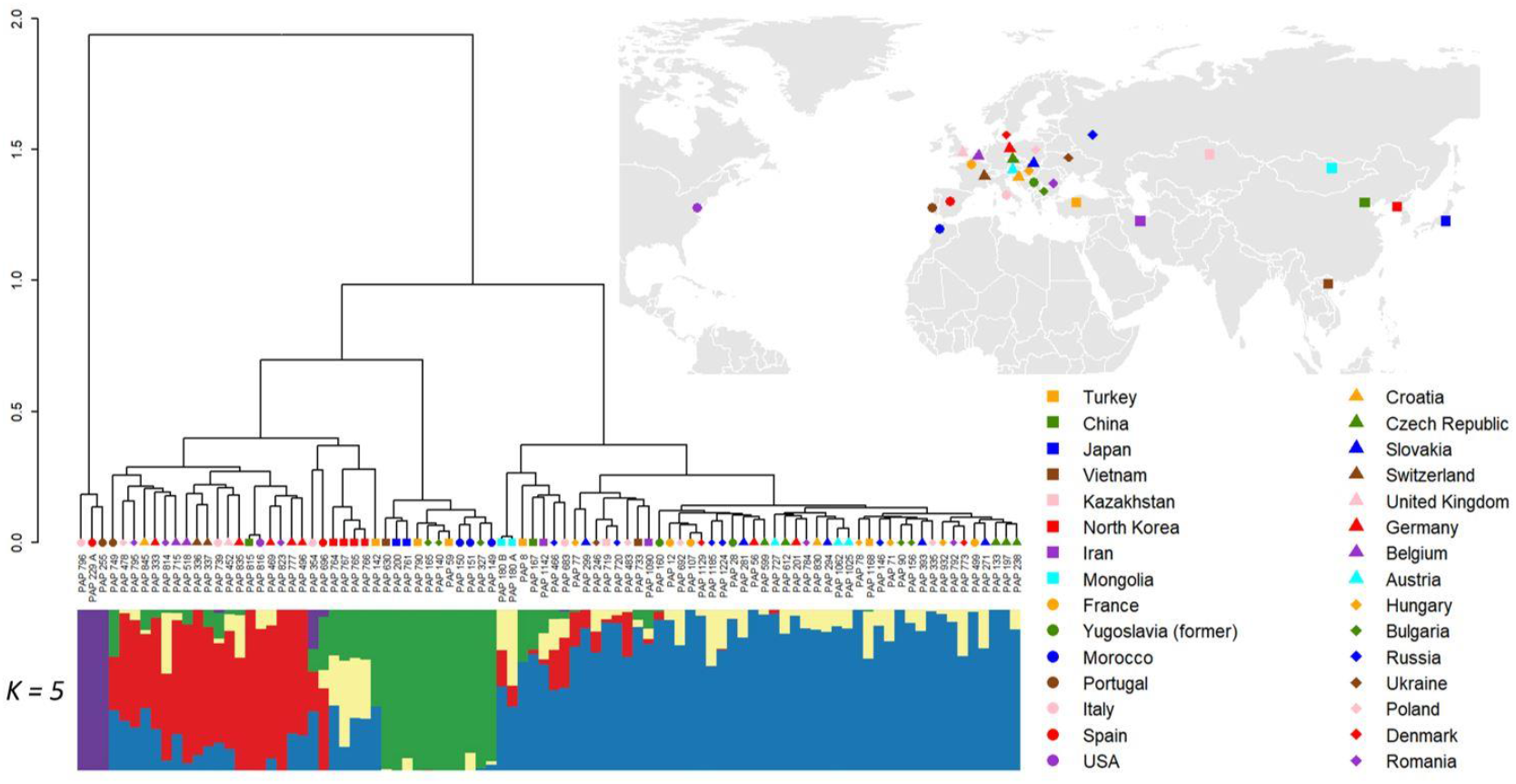
Genetic diversity and structure of 90 *Papaver* accessions. IBS clustering showing the genetic relationship among the accessions (top left panel), a map showing origin of the accessions (top right panel) and bar plots describing the five subpopulations obtained by structure analyses (lower panel). The accessions are sorted according to IBS clusters. Country of origin of the accession is indicated by differently coloured shapes. The map was generated using the R package ggplot2 v3.3.5 (https://ggplot2.tidyverse.org).

### Variability in alkaloid content across genetically distinct poppy accessions

*Papaver* cultivars and accessions exhibit notable variability in the quantity and composition of their alkaloid contents^38,39^. Understanding the genetic basis of this variability would improve knowledge of alkaloid biosynthesis and may provide tools for breeding and synthetic biology. We examined this by quantifying both the major (morphine, codeine, thebaine) and minor (papaverine and opripavine) alkaloids in dry capsules of the 90 accessions that reached full maturity (Fig. **7**). We observed considerable variation among the accessions in alkaloid content and composition (Fig. **7**). Total alkaloid content ranged from 0.125 (PAP 795) to 1.610 (PAP 784) g/100 g DW, whereas morphine content varied between 0.072 (PAP 229A, *P. setigerum*) and 1.416 (PAP 784) g/100 g DW. Codeine and thebaine contents ranged from 0.002 (PAP 795) to 0.342 (PAP 151) and 0.000 to 0.336 (PAP 719) g/100 g DW, respectively (Fig. **7a**). Eight of the accessions analysed (9%) did not have a detectable level of thebaine. Papaverine and oripavine content ranged from 0.000 to 0.077 and 0.102 g/100 g DW, respectively. We identified 18 accessions with undetectable levels of papaverine and six with undetectable oripavine. These results demonstrate that considerable diversity in total alkaloid contents exists in the accessions studied.

**Fig. 7.**
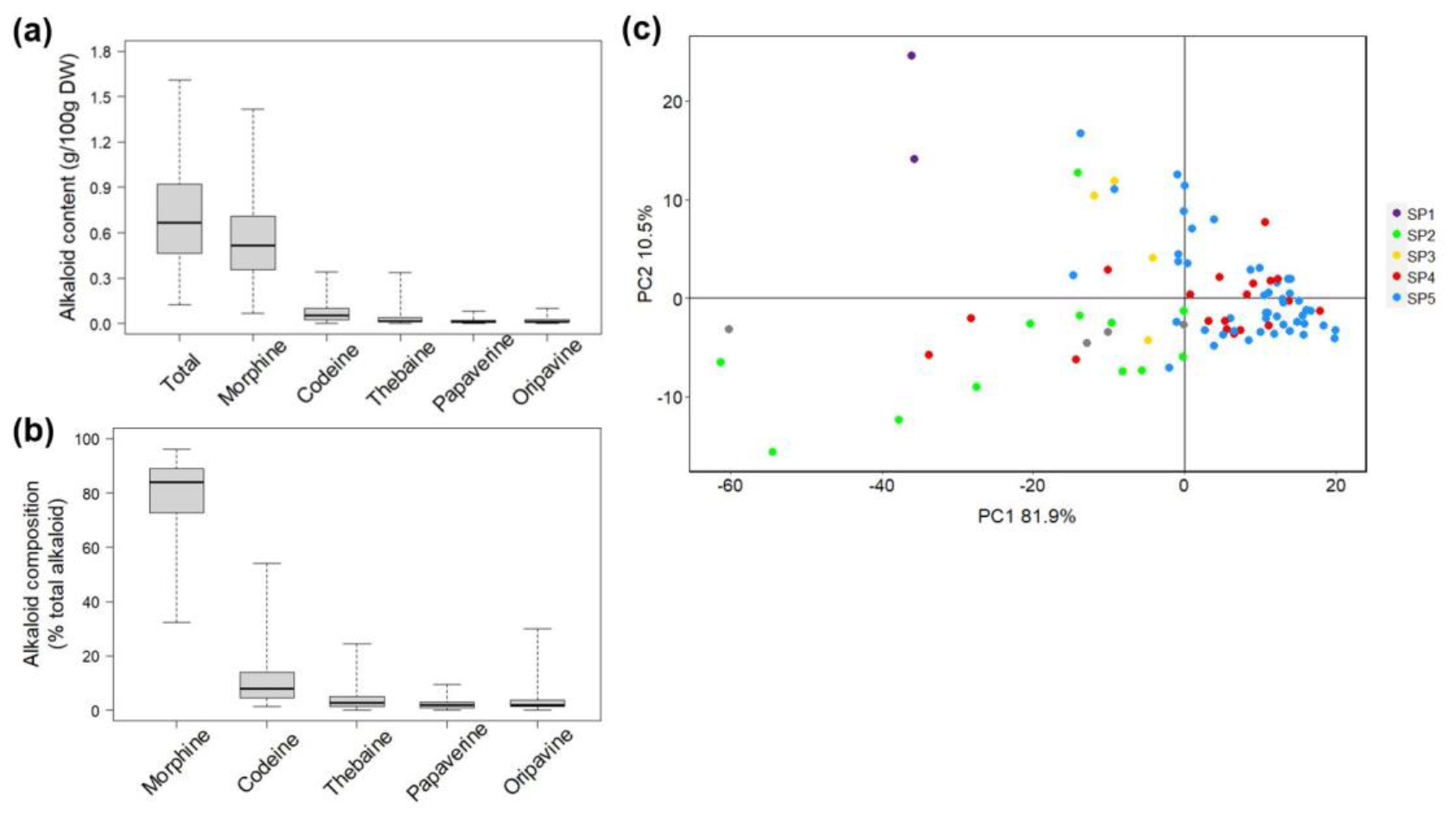
Diversity in alkaloid profiles of *Papaver* accessions. **(a)** The variability in alkaloid content and **(b)** composition of poppy capsules. **(c)** Two-dimensional PCA score plot showing the diversity in alkaloid composition. Axes are labelled with the percentage of variation explained by the two PCs. Colours represent the five subpopulations identified by STRUCTURE analysis (**Fig. 5**). Accessions for which no GBS data was available are indicated with grey dots. The *P. setigerum* accessions (PAP 229A and PAP 255) and the six accession with the highest codeine levels (PAP 151, PAP 152, PAP 150, PAP 200, PAP 739, PAP 149, and PAP 353) are indicated. SP, subpopulation.

The relative abundance of individual alkaloids also varied across accessions. Morphine was the most abundant alkaloid in 85 of the 90 accessions (94.4%), ranging from 32.3% to 96% (of total alkaloids) (Fig. **7b**). Codeine was the most abundant alkaloid (over 50% of total alkaloids) in only three accessions (PAP 150, PAP 151 and PAP 152), though 14 accessions had codeine abundances of 20% (of total alkaloids) or more. Notably, all three high codeine accessions were collected in Morocco. Thebaine, papaverine or oripavine were the most abundant alkaloid in none of the accessions. Thebaine composition ranged from 0.0% to 24.4% (of total alkaloids), with the highest in PAP 719. The proportions of papaverine ranged from 0.0% to 9.4% and oripavine from 0.0% to 29.9% (of total alkaloids).

We tested for relationships between genetic and chemical diversity by applying principal component analysis based upon the alkaloid profiles of the accessions, then compared subpopulation clustering patterns (Fig. **7c** and Fig. **S6**). The accessions of subpopulations 4 and 5 had the most similar alkaloid composition, demonstrated by PCA (PC1 and PC2 accounting for 92.4% of variability, Fig. **7c**). The *P. setigerum* accessions were clearly separated from all others, while accessions of subpopulation 2 had the most diverse and distinct alkaloid composition among the *P. somniferum* accessions. Accessions with the highest codeine proportions, including PAP 150 (53.9%), PAP 151 (52.0%), PAP 152 (50.4%), PAP 200 (42.2%), PAP 739 (35.6%), PAP 149 (33.8%) and PAP 354 (28.7%), were clearly separate from the others (Fig. **7c**). These results suggested a congruent pattern of genetic and chemical diversity of the accessions. However, considering that alkaloid content and composition can be affected by environment, it is important that such data is validated in replicated field experiments^38^. For a crop like poppy that is highly valued for its secondary metabolites, understanding patterns of metabolic variation is important both for conservation and utilization of the available germplasm.

### Phylogenetic relationships of *P. somniferum* accessions

We explored the phylogenetic relationships between *P. somniferum* accessions within the collection by constructing a Neighbor-Net using 49,160 unlinked SNPs^40^. Distinct genetic groups were identified, mostly reflecting the subpopulations identified by the population structure analyses (Fig. **8**). Accessions of *P. somniferum* and *P. setigerum* were clearly separated. Groups 1 and 2 contained all the three and 12 accessions from SP1 and SP2 of the DAPC, respectively. The accessions of SP4 differentiated into two groups, Group 3 and 4, consistent with the STRUCTURE analysis indicating that the highest amount of admixture was in SP4 (Fig. **5b** and **8**). The North Korean accessions formed Group 5, while Group 6 included all 50 accessions of DAPC SP5.

**Fig. 8.**
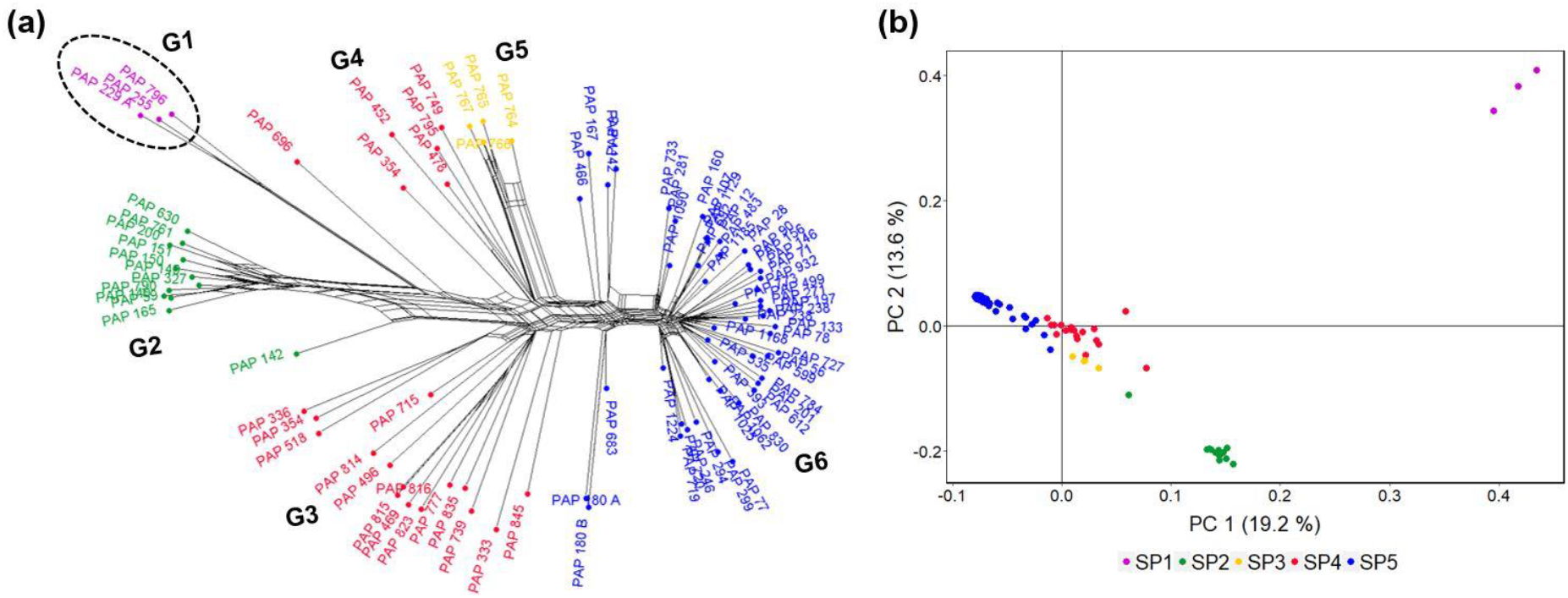
Phylogenetic network analysis of 90 *Papaver* accessions. (**a**) A Neighbor-Net constructed by SplitsTree using 49,160 GBS SNP markers. The six genetic groups identified (G1-G6) are shown. Accessions are labelled with different colours according to the five subpopulations identified by DAPC analysis (**Fig. 5**). The *P. setigerum* accessions, indicated in circle, are differentiated from the *P. somniferum* accessions. (**b**) Principal component (PC) analysis of 90 *Papaver* accessions using 49,166 GBS SNP markers. Axes are labelled with the percentage of variation explained by the two PCs. The five subpopulations (SP1-SP5) identified by DAPC analysis are shown with different colours. SP, subpopulation.

We examined morphological and agronomical traits that could potentially distinguish the various subpopulations and genetic groups. We found that variations in the agronomic-related traits of number of branches per plant and 1000-seed weight were broadly consistent with the subpopulations and phylogenetic groups identified (Fig. **9a**). Branch number ranged from 1.75 (PAP 184) to 11.25 (PAP 229A), and 1000-seed weight from 0.165 g (PAP 255) to 0.815 g (PAP 733). The *P. setigerum* accessions (Group 1) and accessions in Groups 2 to 5, representing accessions of the DAPC SP1 to SP4, were generally highly branching and produced lighter seeds (Fig. **9a**). Contrastingly, accessions in Group 6 (SP5) were less branching and produced heavier seeds. We tested if the association between branch number 1000-seed weight was significant and found a significant negative correlation between the two traits (r=−0.70, *p* < 0.001) (Fig. **9c**). Seed size and branching habit are important characteristics to distinguish *Papaver* species^5,41^. We also observed differences between genetic groups with respect to capsule and seed morphology (Fig. **S4,S5**). The loss of a wind-based seed dispersal mechanism through a transition from poricidal to indehiscent capsules is believed to be among the changes in morphological traits that occurred during poppy domestication^7^. Our study provides a preliminary analysis of this interesting topic, which could be further investigated using larger experiments that incorporate field data.

**Fig. 9.**
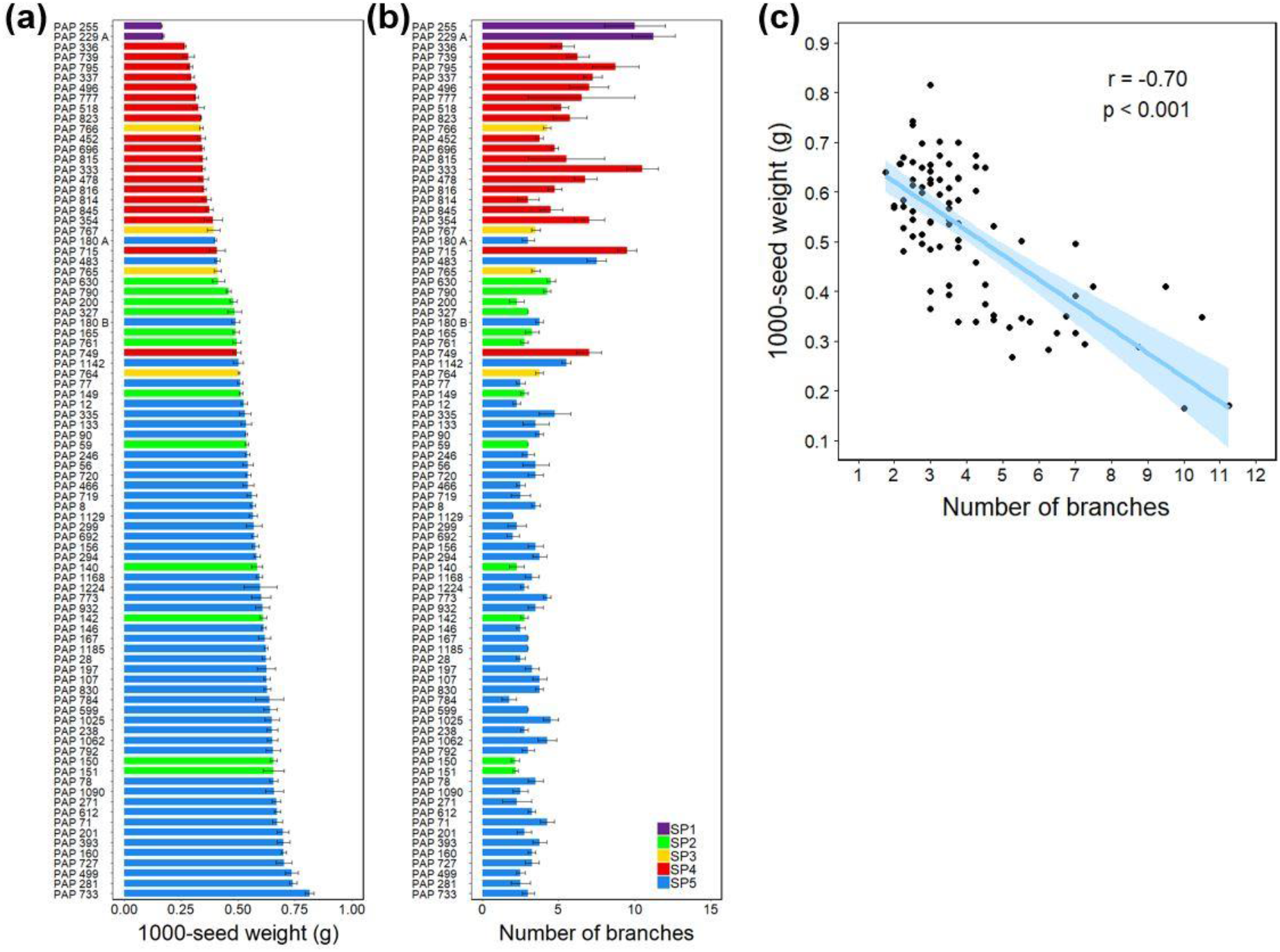
Variations in selected agronomic-related traits in *Papaver* accessions. (**a**) The diversity in number of branches per plant and (**b**) 1000-seed weight among the accession studied. The five subpopulations identified by STRUCTURE and DAPC analyses (**Fig. 5**) are indicated by bars with different colours. Data shown are mean ±SE of three (1000-seed weight) and two to seven (number of branches) biological replicates. SP, subpopulation. (**c**) A scatter plot showing the relationships between number of branches and 1000-seed weight. A linear regression line is shown in blue, and the region shaded in light blue represents the 95% confidence interval. Pearson correlation coefficients (*r*) and their significance level (*p*) are shown.

## Discussion

The poppy germplasm currently available in several genebanks is a rich potential source of useful alleles. Accurate genotyping of this germplasm is crucial to dissect the available genetic diversity. This is an important step for germplasm conservation and can also facilitate the identification and deployment of promising genotypes/alleles for genetic improvement of the crop. With this in mind, we optimized a GBS protocol for poppy and applied GBS-based analysis of SNP markers for the assessment of genetic relationships in diverse poppy accessions from *P. somniferum* and *P. setigerum*. This optimized protocol provides a rapid and cost-effective method for genotyping of unlimited number of accessions.

The current *P. somniferum* genome is a draft with only 81.6% of the sequences assigned to individual chromosomes^27^. However, we were able to identify significantly more markers when using it as a reference for both *P. somniferum* and *P. setigerum* SNP calling than when using a reference-free GBS analysis method (Table **S4**). This result was supported by a high alignment rate (> 97%) and number of private alleles (37,157) identified for the *P. setigerum* accessions (Table **2**; Table **S3**). The *P. somniferum* and *P. setigerum* genomes share considerable homology and have a close phylogenetic relationship, which likely explains these results^34,42–46^. The differing ploidy of *P. setigerum* (tetraploid) and *P. somniferum* (diploid) was a potential limitation of our analyses, but GBS has previously been implemented on species with different ploidy levels^47–51^. However, accurate identification of SNPs from polyploids is still a challenge that requires improvement of existing software packages or development of new ones^52^.

Whole genome sequencing (WGS) is an ideal tool to capture complete information about genome variability including SNP variants. However, the RRS approaches such as GBS are practical and cost-effective methods for simultaneous SNP discovery and genotyping of species with large and complex genomes or when analysing large number of samples. Furthermore, for species like opium poppy where the available germplasm is yet to be fully characterized, GBS can be applied to characterize a large collection from which a core set could be selected to represent the genetic diversity of the entire collection. WGS can then be applied to the accessions in the core collection for further, deep analysis. This is a critical step to efficiently manage the genebank collection and promote its use. The accessions characterized in our study and the associated GBS data can significantly contribute towards this goal.

The phylogenetic and taxonomic relationships of *Papaver* species, particularly that of the cultivated *P. somniferum* and the wild species *P. setigerum*, is still debated^6,29,33,46^. Some treat the two as separate species, while others consider *P. setigerum* as a subspecies of *P. somniferum* based on morphology and alkaloid profiles^29,46^. Our findings confirmed the genetic separation of the accessions from the two named groups (Fig. **3**-**5**). This result was expected, because it has been shown previously that *P. setigerum* is distinct from, but phylogenetically closer to, *P. somniferum* than all other *Papaver* species^45,46^. *P. setigerum* is also considered the putative progenitor of cultivated *P. somniferum*^5,29^. Our result based on genome size analysis supports previous reports that *P. somniferum* is diploid (2n = 2× = 22) and *P. setigerum* tetraploid (2n = 4× = 44) (Fig. **4**)^34,35^. These data do not seem to support the hypothesis of a direct origin of *P. somniferum* from *P. setigerum*. However, they raise interesting questions about the possible relationships between the two species. Considerable homology exists between the two genomes, suggesting they may have a common origin^34^. Interspecific crosses are possible between the two species despite some meiotic abnormalities observed at the F1 generation^34,53^. Notably, a diploid form of *P. setigerum* has been described^29,54^. However, our data is based on only three *P. setigerum* accessions that are all tetraploids. For a detailed analysis of the genetic relationship of these two species, more samples from *P. setigerum*, including both the diploid and tetraploid forms, need to be studied.

Our analysis of population structure and phylogeny generated similar accession groupings. Interestingly, these groups can be broadly identified based on traits such as branching and seed weight (Fig. **9**). Although this finding needs further verification with larger datasets and in replicated experiments including in the field, the data can be a useful input for studies aiming to unravel the domestication history of opium poppy. Domestication traits in poppy are poorly defined, making it difficult to investigate this process particularly in archaeological records. Changes in capsule and seed sizes are believed to be among the domestication-related traits in poppy^7^. Our preliminary observations also suggest capsule indehiscence is a useful trait for consideration in future studies.

The accessions we studied were highly diverse in their alkaloid profiles. Although morphine was the dominant alkaloid in most of the accessions, we also identified accessions with codeine levels of up to 54% (of total alkaloids). Both natural and induced mutants with altered alkaloid profiles have been instrumental to elucidate the molecular mechanisms underlying differences in alkaloid contents in poppy^27,55–58^. The transcriptional regulation of alkaloid biosynthesis in poppy has not been studied in detail. Diverse chemotypes are potential sources of gene expression or enzyme variants with differing activities that affect the alkaloid biosynthesis pathway. These are vital resources to understand the biochemical and genomic regulation of alkaloid biosynthesis. This is also key for breeding commercial lines with specific alkaloid abundances or desired alkaloid profiles. The development of the *top1* mutant, a high-thebaine and high-oripavine variety, was a significant commercial breakthrough that allowed the production of thebaine from morphine-free plants^56^. Thebaine is used for the semi-synthesis of painkillers oxycodone and hydrocodone and the anti-opioid addiction drugs buprenorphine and naltrexone^59^. Similarly, the development of high codeine and morphine-free poppy varieties would allow the direct plant-based production of codeine while preventing the illicit synthesis of the morphine-derived heroin^60^.

Our study demonstrates the utility of GBS for genetic analyses in opium poppy. Many hundreds of poppy germplasm accessions are available from genebanks worldwide. These are an immense potential resource if fully exploited. Application of GBS to the entire poppy germplasm collection, complemented by the currently available draft reference genome sequence of poppy, could drive further studies to unravel the extent of genetic diversity in the species. Although poppy is largely self-pollinating, a considerable degree of outcrossing has been reported^61–63^. Consequently, future genetic diversity studies need to take into consideration intra-accession variability. Based on our preliminary observations, we also suggest that traits such as seed weight, capsule size and dehiscence, and branching be considered in studies of the domestication and phylogenetic relationships of opium poppy and related species. Data generated from such studies would also enable the development a poppy core set, which is an important step towards the selection of suitable parents and development of mapping populations or genetic panels for elucidating the genetic architecture of traits of agronomic and pharmaceutical importance.

## Methods

### Plant materials and morphological characterization

Seeds of 95 *Papaver* accessions from diverse geographical origins were obtained from the global poppy germplasm collection maintained at Leibniz Institute of Plant Genetics and Crop Plant Research (IPK) Genbank in Germany (Table **S1**). Of these, 91 successfully germinated accessions and were used for GBS analysis. We excluded four *P. somniferum* accessions that failed to germinate. The accessions were morphologically diverse materials belonging to the two main *Papaver* species: *P. somniferum* (88 accessions) and *P. setigerum* (3 accessions) (Fig. **1**). Seeds were sown in 200 mm pots using a standard potting mix. Two pots (each a replication) were used per line and plants were thinned to two per pot after germination. Plants were grown in a glasshouse under constant temperature (22 °C day and night) and photoperiod (17 hr/7 hr light/dark). Capsules were harvested for alkaloid profiling at the dry capsule stage.

At full plant maturity data was recorded on number of branches (main stem plus side branches), 1000-seed weight, seed colour and capsule and seed characteristics. Images of capsules were taken under normal room light using a SM-G950U1 digital camera. The seed images were captured using a Leica M205 FA stereomicroscope with a digital camera Leica DFC 420 and Leica Application Suite Software (LAS 4.0, Leica) with 10% light and 50% zoom in condition. To measure seed weight, we first removed debris then counted 1000 clean seeds using a Contador seed counter fitted with feed container no. 3 for fine seeds (Pfeuffer GmbH, Germany). To determine seed colour, images of the seeds were taken using a Cannon EOS-600D camera together with a ColorChecker passport (X-rite) for camera calibration and colour correction. The original Red, Green, Blue (RGB) colour channels of seed digital images were calibrated in Lightroom Classic CC (Adobe Creative Cloud). The RGB values were then converted to Hex colour codes using RGB Color code chart (https://www.rapidtables.com/web/color/RGB_Color.html). The corresponding approximate colours were determined from Hex colour codes using HTML CSS Color Picker (https://www.htmlcsscolor.com/hex/).

### DNA isolation, GBS library preparation and sequencing

For DNA extraction, ~50-100 mg fresh young leaves were harvested in liquid nitrogen and ground to a fine powder using a TissueLyser II system (Qiagen). Genomic DNA was extracted using the CTAB (cetyl trimethylammonium bromide) method^64^ with the following minor modification.

Following grinding of the samples but prior to addition of the extraction buffer, samples were washed using a 0.1 M HEPES (N-2-hydroxyetylpiperazine-N′-ethanesulphonic acid) buffer (pH 8.0) containing 1% polyvinylpyrrolidone, 0.9% L-ascorbic acid, and 2% 2-mercaptoethanol to remove polysaccharides and phenolic compounds. DNA quality was checked by agarose gel electrophoresis and quantified with a NanoDrop spectrophotometer ND-1000 version (Thermo Fisher Scientific, Wilmington, DE, USA). For GBS library construction, the double digest RAD-seq (ddRAD) based library preparation protocol was used^65^. The protocol included the following steps: DNA digestion with two restriction enzymes, ligation of barcoded adapters compatible with restriction sites overhang, size selection of pooled digested-ligated fragments using Blue Pippin, and amplification of library via PCR using indexed primers. For protocol optimization, eight double digested libraries were prepared from a pool of 3 representative samples (PAP 630, PAP 696 and a commercial cultivar) using the restriction enzyme combinations *Pst*I/*Msp*I, *Pst*I/*Mse*I, *Pst*I/*Nla*III, *Pst*I/*Hpy*CH4IV, *Eco*RI/*Msp*I, *Eco*RI/*Mse*I, *Eco*RI/*Nla*III and *Eco*RI/*Hpy*CH4IV. The library generated using *Eco*RI and *Nla*III was sequenced on the Illumina NextSeq500 platform (Illumina, San Diego, CA, USA) with the standard protocol for single-end reads in 150-cycle mid-output mode at the Australian Genome Research Facility (Melbourne, Australia).

### Mapping and SNP calling

Raw sequence data were demultiplexed and sorted using the “process_radtags” function in STACKS v2.41 with the default parameters: “--inline-index” for barcode option and “--renz_1 EcoRI--renz_2 NlaIII” for enzymes option^66^. After trimming barcode sequences, all trimmed reads (150 bp) were checked for quality, and low-quality reads (with quality score of less than 10) and “no RadTag” reads were removed. The filtered reads were aligned to the draft poppy genome sequence retrieved from NCBI (GCA_003573695.1_ASM357369v1) using BWA v0.7.17 and SAMtools v1.9^27,67,68^. SNP calling was performed using the refmap.pl pipeline of STACKS v2.41. All the 91 accessions were treated as a single population. SNP loci that were mapped with reads from less than 90% of the accessions sequenced (--min-samples-per-pop 0.9) were excluded from further analysis. The same parameter was applied for the non-reference-based SNP calling using denovo.pl pipeline of STACKS. The density and distribution of the filtered SNPs across poppy chromosomes were determined using BEDTools v2.26.0 and SnpEff v4.1l^69,70^. To visualize SNP density, the “MVP.report” function with 1Mb non-overlapping window from rMVP package was performed in R studio^71–73^.

### Population structure analysis

The population structure of the accessions was assessed using four methods. First, the accessions were clustered using identity-by-state (IBS) to determine the relationship between accessions based on the proportion of shared alleles between pairs of individuals in PLINK v1.07^74^. Second, for elucidation of the genetic structure and identification of the optimum number of subgroups, we removed the outlier PAP 400 and applied the Bayesian information criterion (BIC) analysis to the remaining 90 accessions. BIC analysis as a nonparametric method was performed with the adegenet package v2.1.3 in R studio based on 131,039 SNP dataset^71,75,76^. The best number of subgroups/subpopulations was determined as the *K*-means corresponding to the lowest BIC score using the “find.clusters” function. To determine the relationship between the subpopulations, we carried out discriminant analysis of principal components (DAPC). A cross validation function “Xval.dapc” was used to determine the optimal number of PCs to be retained.

Third, to investigate the population structure in detail, the admixture within the accessions was determined using Bayesian clustering based on a Bayesian Markov Chain Monte Carlo model (MCMC) implemented in STRUCTURE v2.3.4^37^. With the assumption that SNPs at the same RAD locus are linked, --write_single_snp flag (in STACKS) was applied to ensure that only one SNP per RAD locus was used for STRUCTURE analysis. To determine the most likely number of subpopulations (clusters), four independent runs with 500,000 iterations and a 150,000-step burn-in period were performed for each *K* from 1 to 10. The output was obtained by structureHarvester v0.6.93 using the maximum estimated log-likelihood [log(P(X|K)] model and the highest ∆K in Evanno method^37,77,78^. After determining the most probable *K* values, ten runs of 500,000 iterations followed by a 150,000 step burn-in were performed using STRUCTURE for each *K*. Additionally, for each optimal *K*, CLUMPP was used to generate individual and population Q matrices from the membership coefficient matrices of the ten replicates obtained from STRUCTURE^79^. Bar plots were generated using DISTRUCT software^80^. Forth, principal component analysis (PCA) was conducted using PLINK v1.07 and plotted using ggplot2 package in R studio ^71,74,81^.

### Genetic diversity and differentiation

Common measures of genetic diversity including private allele number (AP), percentage of polymorphic loci (%Poly), observed and expected heterozygosity (*H_O_* and *H_E_*), nucleotide diversity (*π*) and inbreeding coefficient (*F_IS_*) were calculated for the five subpopulations using the “populations” function in STACKS v2.41^66^. The genetic differentiation between the subpopulations was calculated based on pairwise population differentiation (*F*_ST_) values from GENODIVE v3.0582. Significance levels (α = 0.05) of the *F*_ST_ values were determined by running 999 permutations and assessing this against a Bonferroni-adjusted *P*-value to account for multiple testing. The correlation matrix was visualized using the corrplot package in R studio^73,83^. To determine the distribution of genetic variation, analysis of molecular variance (AMOVA) was performed using GENODIVE v3.05^82^. Significance level was tested using 999 permutations.

### Phylogenetic analysis

To explore the phylogenetic relationships between *P. somniferum* accessions, we constructed a phylogenetic network using 49,160 unlinked SNPs. The SNPs were exported in PHYLIP format (--phylip-var) from the “populations” function in STACKS v2.41 to the SplitTree4 software^40^. The split network was created with the “uncorrectedP” method, which ignores ambiguous sites, and visualized using “Neightbornet” network. One thousand bootstrap values were used.

### Alkaloid profiling

Alkaloid content of 90 accessions was measured according the protocol used by Dittbrenner and colleagues^38^. Only accessions that produced capsules samples sufficient for analysis were included. Alkaloids were also analysed in four accessions that had failed to germinate during the first trial and subsequently lacked GBS data (Table **S1**). For testing the relationship between alkaloid and genetic diversity, we conducted principal component analysis (PCA) using TASSELv5.2.73 and plotted the graph using ggplot2 package in R studio^73,81,84^.

### Genome size and ploidy analysis

The genome size and ploidy level of selected accessions was estimated using flow cytometry analysis of propidium iodide (PI)-stained nuclei isolated from poppy leaves. Nuclei of tomato (*Solanum lycopersicum*), which has genome size of ~900 Mb^85^, were simultaneously isolated from leaves, stained and analysed with poppy nuclei as an internal reference standard. Nuclei were isolated with the Galbraith lysis buffer using a modified protocol from Gutzat and Scheid as follows: (1) Chopping fresh young leaf tissue (0.5 g) using double-sided razor blades in 2 mL ice-cold lysis buffer, (2) filtering the homogenate through a 40 μm nylon mesh, (3) adding 2.5 μL of RNase (10 mg/mL) to 500 μL filtered homogenate, then incubating on ice for 10 minutes, (4) centrifugation at 400 g for 3 minutes, removing the supernatant and resuspending the pellet gently in 1 mL lysis buffer, then incubating on ice for 15 minutes, and (5) filtering the homogenate again through a 40 μm nylon mesh^86,87^. For nuclei staining, 25 μL PI (1 mg/ml) was added into 500 μL nuclei solution to get a final concentration of 50 μg/mL. Samples were screened on a CytoFLEX S flow cytometer (Beckman Coulter, Brea, CA, USA). The PerCP-A fluorescence intensity of G1 and G2 phase cells of internal standard and samples was used to estimate genome size and ploidy level of the samples.

## Supporting information

All supplements

## Acknowledgments

UTVH received a PhD scholarship from La Trobe University Graduate Research School. Work in the Lewsey lab is funded by the Australian Research Council Industrial Transformation Hub in Medicinal Agriculture (IH180100006) and a Commonwealth Scientific and Industrial Research Organisation SIEF STEM+ Fellowship with Palla Pharma Ltd. We thank La Trobe’s Bioimaging Platform for support with genome size analysis.

## Authors’ contributions

MTO, MGL, UVTH designed the study. UVTH, MTO, CRO, ARA conducted the experiments. UVTH, MTO, BH analysed the data. CRO, ARA provided research materials. UVTH, MTO, MGL interpreted the results and wrote the paper. All co-authors read and approved the final manuscript.

## Competing interests

UVTH, MTO, BH and MGL declare no competing interests. CRO and ARA are employees of Palla Pharma Ltd.

## Ethical standards

Seeds of all poppy accessions were obtained from a public seedbank and transferred in accordance with international legislation. All experimental research in this study was conducted in compliance with the relevant institutional, national, and international guidelines and legislation.

## Notes

### Competing Interest Statement

Chris R. Okey and Artur R. Abreu are employees of Palla Pharma Ltd.

### Summary of Updates

Main text and supplemental files updated.

