## Supplementary material for "Insights into opium poppy (*Papaver* spp.) genetic diversity from genotyping-by-sequencing analysis": All supplements

**Table S1.** List of the *Papaver* accessions include in this study. All accessions were sourced from the poppy germplasm collections at the Leibniz Institute of Plant Genetics and Crop Plant Research (IPK), Germany.

| No. | Acc ID | Species | Origin | Common name |
|---|---|---|---|---|
| 1 | PAP 8 | *P. somniferum* | Turkey | |
| 2 | PAP 12 | *P. somniferum* | France | Tourneur-Freres-Typ D |
| 3 | PAP 28 | *P. somniferum* | Yugoslavia (former) | |
| 4 | PAP 56 | *P. somniferum* | Germany | |
| 5 | PAP 59 | *P. somniferum* | Turkey | |
| 6 | PAP 71 | *P. somniferum* | Hungary | Pulawski Niebieski |
| 7 | PAP 77 | *P. somniferum* | Hungary | Pitvarositf |
| 8 | PAP 78 | *P. somniferum* | Hungary | Fertodi Soktoku Kek |
| 9 | PAP 90 | *P. somniferum* | Bulgaria | Voronezskij 1042 |
| 10 | PAP 107 | *P. somniferum* | France | Flora II |
| 11 | PAP 133 | *P. somniferum* | Czechoslovakia | Erbachshofsky |
| 12 | PAP 140 | *P. somniferum* | Bulgaria | |
| 13 | PAP 142 | *P. somniferum* | Turkey | |
| 14 | PAP 146 | *P. somniferum* | Russia | Altaec |
| 15 | PAP 149 | *P. somniferum* | Morocco | violet plat |
| 16 | PAP 150 | *P. somniferum* | Morocco | coralla v. majore |
| 17 | PAP 151 | *P. somniferum* | Morocco | blanc large |
| 18 | PAP 152[a] | *P. somniferum* | Morocco | blanc carre |
| 19 | PAP 156 | *P. somniferum* | Bulgaria | Tatarstan I |
| 20 | PAP 160 | *P. somniferum* | Yugoslavia (former) | |
| 21 | PAP 165 | *P. somniferum* | Bulgaria | Fertodi Zarttoku Kek |
| 22 | PAP 167 | *P. somniferum* | China | |
| 23 | PAP 197 | *P. somniferum* | Czechoslovakia | Susicky Cervenosemenny |
| 24 | PAP 200 | *P. somniferum* | Japan | Ikkanshu |
| 25 | PAP 201 | *P. somniferum* | Germany | Daubaer Silbergrauer |
| 26 | PAP 238 | *P. somniferum* | Czechoslovakia | Mahndorfer |
| 27 | PAP 246 | *P. somniferum* | Ukraine | Lubenskij 6 |
| 28 | PAP 255 | *P. setigerum* | Portugal | |
| 29 | PAP 271 | *P. somniferum* | Slovakia | |
| 30 | PAP 281 | *P. somniferum* | Slovakia | |
| 31 | PAP 294 | *P. somniferum* | Slovakia | |
| 32 | PAP 299 | *P. somniferum* | Slovakia | |
| 33 | PAP 327 | *P. somniferum* | Bulgaria | Sinjocaftjast P 360 |
| 34 | PAP 333 | *P. somniferum* | Germany | |
| 35 | PAP 335 | *P. somniferum* | Poland | |
| 36 | PAP 336 | *P. somniferum* | Switzerland | |
| 37 | PAP 337 | *P. somniferum* | Switzerland | |
| 38 | PAP 354 | *P. somniferum* | Italy | |
| 39 | PAP 393 | *P. somniferum* | Slovakia | |
| 40 | PAP 400[b] | *P. somniferum* | Spain | |
| 41 | PAP 452 | *P. somniferum* | United Kingdom | Pink Chiffan |
| 42 | PAP 466 | *P. somniferum* | Russia | Daneborg |
| 43 | PAP 469[b] | *P. somniferum* | Germany | |
| 44 | PAP 478 | *P. somniferum* | Poland | |
| 45 | PAP 483 | *P. somniferum* | Poland | |
| 46 | PAP 496 | *P. somniferum* | Germany | |
| 47 | PAP 499 | *P. somniferum* | France | |

| | | | | |
|---|---|---|---|---|
| 48 | PAP 518 | *P. somniferum* | Belgium | |
| 49 | PAP 599 | *P. somniferum* | Czechoslovakia | Hatvani |
| 50 | PAP 612 | *P. somniferum* | Czechoslovakia | Selecty Stribrosedy |
| 51 | PAP 630 | *P. somniferum* | Vietnam | |
| 52 | PAP 683[b] | *P. somniferum* | Italy | |
| 53 | PAP 692 | *P. somniferum* | Poland | |
| 54 | PAP 696 | *P. somniferum* | Spain | |
| 55 | PAP 715 | *P. somniferum* | Belgium | |
| 56 | PAP 719 | *P. somniferum* | Kazakhstan | |
| 57 | PAP 720 | *P. somniferum* | Russia | |
| 58 | PAP 727 | *P. somniferum* | Austria | |
| 59 | PAP 733 | *P. somniferum* | Vietnam | |
| 60 | PAP 739 | *P. somniferum* | Italy | |
| 61 | PAP 747[a] | *P. somniferum* | Portugal | |
| 62 | PAP 749 | *P. somniferum* | Portugal | |
| 63 | PAP 761 | *P. somniferum* | Japan | Ikkan |
| 64 | PAP 764 | *P. somniferum* | North Korea | |
| 65 | PAP 765 | *P. somniferum* | North Korea | |
| 66 | PAP 766 | *P. somniferum* | North Korea | |
| 67 | PAP 767 | *P. somniferum* | North Korea | |
| 68 | PAP 768[a] | *P. somniferum* | North Korea | |
| 69 | PAP 773 | *P. somniferum* | Denmark | Parmo |
| 70 | PAP 777 | *P. somniferum* | Germany | |
| 71 | PAP 784 | *P. somniferum* | Romania | |
| 72 | PAP 790 | *P. somniferum* | Turkey | |
| 73 | PAP 792 | *P. somniferum* | Romania | |
| 74 | PAP 795 | *P. somniferum* | Romania | |
| 75 | PAP 796[b] | *P. setigerum* | Italy | |
| 76 | PAP 815 | *P. somniferum* | China | |
| 77 | PAP 814 | *P. somniferum* | Romania | |
| 78 | PAP 816 | *P. somniferum* | USA | Pink Single Poppy |
| 79 | PAP 823[b] | *P. somniferum* | Romania | mac |
| 80 | PAP 827[a] | *P. somniferum* | Croatia | mak (kroatischer Name fur Mohn) |
| 81 | PAP 830 | *P. somniferum* | Croatia | mak |
| 82 | PAP 835 | *P. somniferum* | Germany | |
| 83 | PAP 845 | *P. somniferum* | Croatia | |
| 84 | PAP 932 | *P. somniferum* | Hungary | KASSA |
| 85 | PAP 1025 | *P. somniferum* | Austria | |
| 86 | PAP 1062 | *P. somniferum* | Austria | |
| 87 | PAP 1090 | *P. somniferum* | Iran | MOHN AUS PERSIEN II |
| 88 | PAP 1129 | *P. somniferum* | Denmark | LORI |
| 89 | PAP 1142 | *P. somniferum* | Iran | MOHN AUS PERSIEN |
| 90 | PAP 1168 | *P. somniferum* | Turkey | NYAZI MUTAFT |
| 91 | PAP 1185 | *P. somniferum* | Russia | VILAR-2 |
| 92 | PAP 1224 | *P. somniferum* | Russia | MAJAK |
| 93 | PAP 180 A | *P. somniferum* | Mongolia | |
| 94 | PAP 180 B | *P. somniferum* | Mongolia | |
| 95 | PAP 229 A | *P. setigerum* | Spain | |

[a]Accessions not included for GBS analysis.
[b]Accessions not included for alkaloid profiling.

**Table S2.** Summary statistical of data generated from GBS of 91 *Papaver* accessions.

| **Steps** | **Statistics** | **Total size (Gb)** |
|---|---|---|
| **Data generation and alignment** | | |
| Raw reads | 103,802,122 | 15.57 |
| Average number of raw reads per sample | 1,140,683 | |
| Number of low quality (<10) reads removed | 89,099 | |
| Number of "no Radtag" removed | 111,744 | |
| Number of filtered reads retained | 103,601,279 | 15.54 |
| Average number of filtered read per sample | 1,134,476 | |
| Average mapping rate per sample | 99.78% | |
| **SNP calling** | | |
| Total loci | 601,567 | |
| Retained loci | 76,407 | |
| Total SNPs | 165,363 | |

**Table S3**. Summary of total raw GBS reads generated, reads retained after filtering and alignment rate for all the 91 *Papaver* accessions sequenced.

| No | Accession ID | Raw reads | Filtered reads | Alignment rate (% ) |
|---|---|---|---|---|
| 1 | PAP 8 | 1,372,600 | 1,368,192 | 99.83 |
| 2 | PAP 12 | 1,121,416 | 1,119,700 | 99.87 |
| 3 | PAP 28 | 1,281,905 | 1,280,107 | 99.87 |
| 4 | PAP 56 | 1,397,053 | 1,394,964 | 99.86 |
| 5 | PAP 59 | 1,282,005 | 1,280,480 | 99.85 |
| 6 | PAP 71 | 914,503 | 913,099 | 99.87 |
| 7 | PAP 77 | 1,419,364 | 1,414,747 | 99.87 |
| 8 | PAP 78 | ,661,836 | 1,659,405 | 99.89 |
| 9 | PAP 90 | 1,451,089 | 1,449,191 | 99.89 |
| 10 | PAP 107 | 1,570,650 | 1,568,447 | 99.88 |
| 11 | PAP 133 | 1,392,292 | 1,390,512 | 99.88 |
| 12 | PAP 140 | 1,270,996 | 1,265,976 | 99.71 |
| 13 | PAP 142 | 1,367,621 | 1,363,011 | 99.83 |
| 14 | PAP 146 | 1,109,457 | 1,107,676 | 99.87 |
| 15 | PAP 149 | 1,042,857 | 1,041,270 | 99.81 |
| 16 | PAP 150 | 1,247,034 | 1,245,198 | 99.82 |
| 17 | PAP 151 | 967,586 | 966,278 | 99.82 |
| 18 | PAP 156 | 1,109,777 | 1,108,345 | 99.87 |
| 19 | PAP 160 | 1,512,681 | 1,507,223 | 99.86 |
| 20 | PAP 165 | 1,435,248 | 1,433,276 | 99.84 |
| 21 | PAP 167 | 1,580,845 | 1,578,329 | 99.86 |
| 22 | PAP 197 | 1,411,310 | 1,409,104 | 99.88 |
| 23 | PAP 200 | 1,195,684 | 1,193,613 | 99.83 |
| 24 | PAP 201 | 1,291,436 | 1,289,841 | 99.88 |
| 25 | PAP 238 | 1,089,293 | 1,086,422 | 99.88 |
| 26 | PAP 246 | 1,056,573 | 1,055,252 | 99.87 |
| 27 | PAP 255 | 1,188,660 | 1,186,821 | 97.31 |
| 28 | PAP 271 | 810,856 | 809,801 | 99.87 |
| 29 | PAP 281 | 946,870 | 945,700 | 99.87 |
| 30 | PAP 294 | 1,031,744 | 1,030,481 | 99.89 |
| 31 | PAP 299 | 1,594,481 | 1,590,483 | 99.89 |
| 32 | PAP 327 | 1,289,790 | 1,288,211 | 99.85 |
| 33 | PAP 333 | 1,350,891 | 1,348,766 | 99.87 |
| 34 | PAP 335 | 1,358,850 | 1,357,091 | 99.89 |
| 35 | PAP 336 | 1,285,253 | 1,283,741 | 99.84 |
| 36 | PAP 337 | 1,334,204 | 1,332,525 | 99.84 |
| 37 | PAP 354 | 1,241,611 | 1,239,498 | 99.83 |
| 38 | PAP 393 | 987,723 | 986,496 | 99.88 |
| 39 | PAP 400 | 985,611 | 983,763 | 99.68 |
| 40 | PAP 452 | 1,093,853 | 1,091,522 | 99.85 |
| 41 | PAP 466 | 981,975 | 980,759 | 99.86 |
| 42 | PAP 469 | 1,035,352 | 1,034,091 | 99.85 |
| 43 | PAP 478 | 1,166,478 | 1,164,390 | 99.87 |
| 44 | PAP 483 | 1,237,596 | 1,235,857 | 99.88 |
| 45 | PAP 496 | 1,185,615 | 1,183,879 | 99.86 |
| 46 | PAP 499 | 1,199,747 | 1,197,206 | 99.88 |
| 47 | PAP 518 | 1,058,117 | 1,056,776 | 99.84 |
| 48 | PAP 599 | 1,165,399 | 1,163,971 | 99.89 |

| | | | | |
|---|---|---|---|---|
| 49 | PAP 612 | 1,039,595 | 1,030,058 | 99.89 |
| 50 | PAP 630 | 1,238,549 | 1,236,340 | 99.83 |
| 51 | PAP 683 | 1,067,715 | 1,066,161 | 99.86 |
| 52 | PAP 692 | 847,072 | 845,900 | 99.87 |
| 53 | PAP 696 | 1,009,504 | 1,007,944 | 99.83 |
| 54 | PAP 715 | 1,271,977 | 1,270,102 | 99.87 |
| 55 | PAP 719 | 1,286,815 | 1,275,048 | 99.88 |
| 56 | PAP 720 | 1,310,790 | 1,308,355 | 99.87 |
| 57 | PAP 727 | 1,227,385 | 1,225,557 | 99.88 |
| 58 | PAP 733 | 1,123,907 | 1,122,508 | 99.88 |
| 59 | PAP 739 | 1,141,361 | 1,139,632 | 99.84 |
| 60 | PAP 749 | 1,363,536 | 1,361,513 | 99.87 |
| 61 | PAP 761 | 941,915 | 935,642 | 99.83 |
| 62 | PAP 764 | 883,071 | 881,568 | 99.86 |
| 63 | PAP 765 | 893,214 | 892,046 | 99.86 |
| 64 | PAP 766 | 954,639 | 953,144 | 99.87 |
| 65 | PAP 767 | 775,540 | 774,621 | 99.86 |
| 66 | PAP 773 | 797,495 | 796,467 | 99.87 |
| 67 | PAP 777 | 1,447,886 | 1,441,337 | 99.87 |
| 68 | PAP 784 | 1,056,203 | 1,054,466 | 99.88 |
| 69 | PAP 790 | 1,120,950 | 1,119,496 | 99.71 |
| 70 | PAP 792 | 944,459 | 942,565 | 99.88 |
| 71 | PAP 795 | 943,086 | 941,925 | 99.86 |
| 72 | PAP 796 | 1,042,448 | 1,041,158 | 97.51 |
| 73 | PAP 815 | 1,174,597 | 1,170,622 | 99.83 |
| 74 | PAP 814 | 751,683 | 750,690 | 99.84 |
| 75 | PAP 816 | 963,468 | 962,008 | 99.84 |
| 76 | PAP 823 | 1,082,911 | 1,081,699 | 99.84 |
| 77 | PAP 830 | 1,037,429 | 1,036,061 | 99.87 |
| 78 | PAP 835 | 834,364 | 833,384 | 99.86 |
| 79 | PAP 845 | 1,224,644 | 1,220,282 | 99.87 |
| 80 | PAP 932 | 1,178,108 | 1,176,590 | 99.90 |
| 81 | PAP 1025 | 1,028,666 | 1,027,042 | 99.89 |
| 82 | PAP 1062 | 1,000,886 | 999,698 | 99.88 |
| 83 | PAP 1090 | 1,185,619 | 1,184,141 | 99.88 |
| 84 | PAP 1129 | 1,148,524 | 1,146,997 | 99.89 |
| 85 | PAP 1142 | 992,301 | 989,360 | 99.86 |
| 86 | PAP 1168 | 803,204 | 802,144 | 99.89 |
| 87 | PAP 1185 | 772,148 | 771,234 | 99.89 |
| 88 | PAP 1224 | 888,063 | 887,026 | 99.88 |
| 89 | PAP 180 A | 718,216 | 717,236 | 99.86 |
| 90 | PAP 180 B | 1,049,972 | 1,048,685 | 99.86 |
| 91 | PAP 229 A | 1,154,420 | 1,151,341 | 97.62 |

**Table S4.** Comparison of the number of single nucleotide polymorphisms (SNPs) identified by the reference- and nonreference-based GBS pipelines for SNP calling using 91 opium poppy accessions.

| | Number of SNPs called | |
|---|---|---|
| **-r** | **Reference-free (*de novo*)** | **Reference-based** |
| 0 | 224,940 | 513,165 |
| 0.5 | 77,820 | 380,676 |
| 0.6 | 47,836 | 339,763 |
| 0.7 | 21,328 | 293,910 |
| 0.8 | 7,650 | 244,382 |
| 0.9 | 1,835 | 165,363 |

-r, minimum percentage of individuals in a population required to process a locus for that population. Both analyses were conducted in STACKS.

**Table S5.** Potential effects of the single nucleotide polymorphisms (SNPs) identified by GBS on the *P. somniferum* genome (annotation GCA_003573695.1_ASM357369v1) analysed using SnpEff v4.11.

| **Number of effects** | **165,786** | **100.00%** |
|---|---|---|
| Genic regions | 16,250 | 9.80% |
| Exon | 5,365 | 3.24% |
| Intron | 7,546 | 4.55% |
| 5’ UTR | 1,236 | 0.75% |
| Splice site donor | 26 | 0.02% |
| Splice site region | 346 | 0.21% |
| Splice site acceptor | 20 | 0.01% |
| Transcript | 8 | 0.01% |
| 3’ UTR | 1,703 | 1.03% |
| Intergenic regions | 149,536 | 90.20% |

**Table S6.** Summary of seed colour of the *Papaver* accessions included in this study. The RGB values were estimated from raw pictures using ColorChecker (X-rite) in Adobe Lightroom Classic CC software. The RGB values were converred to Hex colour codes with RGB Color code chart, and the corresponsding approximate colours were determined from Hex colour codes using HTML CSS Color Picker.

| No. | Accession ID | RGB value | | | HEX colour code | Approximate colour |
|---|---|---|---|---|---|---|
| | | R | G | B | | |
| 1 | PAP 8 | 109 | 98 | 85 | #6D6255 | Makara |
| 2 | PAP 12 | 58 | 55 | 58 | #3A373A | Fuscous Grey |
| 3 | PAP 28 | 74 | 67 | 65 | #4A4341 | Crater Brown |
| 4 | PAP 56 | 54 | 53 | 56 | #363538 | Martinique |
| 5 | PAP 59 | 162 | 144 | 112 | #A29070 | Pale Oyster |
| 6 | PAP 71 | 120 | 121 | 122 | #787979 | Rolling Stone |
| 7 | PAP 77 | 88 | 80 | 69 | #585045 | Mondo |
| 8 | PAP 78 | 56 | 59 | 68 | #383B44 | Vulcan |
| 9 | PAP 90 | 60 | 62 | 67 | #3C3E43 | Vulcan |
| 10 | PAP 107 | 109 | 104 | 102 | #6D6866 | Russett |
| 11 | PAP 133 | 46 | 48 | 55 | #2E3037 | Ebony |
| 12 | PAP 140 | 166 | 152 | 127 | #A6987F | Bronco |
| 13 | PAP 142 | 92 | 75 | 68 | #5C4B44 | Saddle |
| 14 | PAP 146 | 56 | 58 | 62 | #383A3E | Vulcan |
| 15 | PAP 149 | 53 | 53 | 55 | #353537 | Revolver |
| 16 | PAP 150 | 97 | 82 | 62 | #61523E | Judge Grey |
| 17 | PAP 151 | 169 | 151 | 125 | #A9977D | Bronco |
| 18 | PAP 152 | 164 | 147 | 116 | #A49374 | Pale Oyster |
| 19 | PAP 156 | 61 | 58 | 63 | #353537 | Revolver |
| 20 | PAP 160 | 102 | 103 | 103 | #666767 | Nevada |
| 21 | PAP 165 | 66 | 68 | 70 | #424446 | Arsenic |
| 22 | PAP 167 | 62 | 64 | 73 | #3E4049 | Blue Zodiac |
| 23 | PAP 197 | 109 | 97 | 97 | #6D6161 | Zambezi |
| 24 | PAP 200 | 172 | 160 | 135 | #ACA087 | Napa |
| 25 | PAP 201 | 121 | 116 | 112 | #797470 | Sand Dune |
| 26 | PAP 238 | 59 | 62 | 68 | #3B3E44 | Vulcan |
| 27 | PAP 246 | 46 | 43 | 41 | #2E2B29 | Bokara Grey |
| 28 | PAP 255 | 27 | 26 | 27 | #1B1A1B | Mardi Gras |
| 29 | PAP 271 | 61 | 63 | 68 | #3D3F44 | Vulcan |
| 30 | PAP 281 | 63 | 66 | 72 | #3F4248 | Steel Grey |
| 31 | PAP 294 | 97 | 90 | 79 | #615A4F | Makara |
| 32 | PAP 299 | 67 | 62 | 60 | #433E3C | Paco |
| 33 | PAP 327 | 61 | 61 | 66 | #3D3D42 | Payne's Grey |
| 34 | PAP 333 | 30 | 27 | 26 | #1E1B1A | Wood Bark |
| 35 | PAP 335 | 120 | 116 | 114 | #787472 | Hurricane |
| 36 | PAP 336 | 32 | 29 | 25 | #201D19 | Bokara Grey |
| 37 | PAP 337 | 37 | 32 | 27 | #25201B | Bokara Grey |
| 38 | PAP 354 | 43 | 36 | 28 | #2B241C | Black Magic |
| 39 | PAP 393 | 129 | 123 | 117 | #817B75 | Americano |
| 40 | PAP 400 | 43 | 38 | 36 | #2B2624 | Wood Bark |
| 41 | PAP 452 | 33 | 30 | 27 | #211E1B | Bokara Grey |
| 42 | PAP 466 | 139 | 120 | 85 | #8B7855 | Cement |
| 43 | PAP 469 | 41 | 35 | 30 | #29231E | Bokara Grey |
| 44 | PAP 478 | 40 | 37 | 36 | #282524 | Wood Bark |

| | | | | | | |
|---|---|---|---|---|---|---|
| 45 | PAP 483 | 46 | 39 | 28 | #2E271C | Black Magic |
| 46 | PAP 496 | 35 | 30 | 26 | #231E1A | Bokara Grey |
| 47 | PAP 499 | 103 | 104 | 103 | #676867 | Finlandia |
| 48 | PAP 518 | 36 | 31 | 28 | #241F1C | Bokara Grey |
| 49 | PAP 599 | 102 | 102 | 108 | #66666C | Comet |
| 50 | PAP 612 | 98 | 88 | 64 | #625840 | Judge Grey |
| 51 | PAP 630 | 121 | 116 | 110 | #79746E | Sand Dune |
| 52 | PAP 683 | 41 | 38 | 39 | #292627 | Aubergine |
| 53 | PAP 692 | 56 | 57 | 64 | #383940 | Payne's Grey |
| 54 | PAP 696 | 33 | 29 | 24 | #211D18 | Bokara Grey |
| 55 | PAP 715 | 48 | 40 | 34 | #302822 | Cocoa Brown |
| 56 | PAP 719 | 59 | 58 | 57 | #3B3A39 | Kilamanjaro |
| 57 | PAP 720 | 55 | 55 | 54 | #373736 | El Paso |
| 58 | PAP 727 | 109 | 102 | 100 | #6D6664 | Russett |
| 59 | PAP 733 | 51 | 50 | 57 | #333239 | Revolver |
| 60 | PAP 739 | 38 | 34 | 29 | #26221D | Bokara Grey |
| 61 | PAP 747 | 33 | 30 | 26 | #211E1A | Bokara Grey |
| 62 | PAP 749 | 116 | 99 | 91 | #74635B | Dorado |
| 63 | PAP 761 | 169 | 153 | 125 | #A9997D | Bronco |
| 64 | PAP 764 | 110 | 104 | 100 | #6E6864 | Dorado |
| 65 | PAP 765 | 87 | 80 | 67 | #575043 | Mondo |
| 66 | PAP 766 | 83 | 77 | 62 | #534D3E | Panda |
| 67 | PAP 767 | 87 | 79 | 61 | #574F3D | Panda |
| 68 | PAP 768 | 159 | 137 | 112 | #9F8970 | Mongoose |
| 69 | PAP 773 | 59 | 63 | 72 | #3B3F48 | Cloud Burst |
| 70 | PAP 777 | 36 | 32 | 29 | #24201D | Bokara Grey |
| 71 | PAP 784 | 119 | 118 | 119 | #777677 | Old Lavender |
| 72 | PAP 790 | 60 | 58 | 63 | #3C3A3F | Martinique |
| 73 | PAP 792 | 114 | 114 | 114 | #727272 | Dim Gray |
| 74 | PAP 795 | 38 | 34 | 27 | #26221B | Black Magic |
| 75 | PAP 796 | 32 | 29 | 28 | #201D1C | Wood Bark |
| 76 | PAP 815 | 49 | 41 | 29 | #31291D | Black Magic |
| 77 | PAP 814 | 43 | 36 | 27 | #2B241B | Black Magic |
| 78 | PAP 816 | 39 | 36 | 29 | #27241D | Maire |
| 79 | PAP 823 | 37 | 34 | 33 | #252221 | Wood Bark |
| 80 | PAP 827 | 57 | 53 | 52 | #393534 | Gondola |
| 81 | PAP 830 | 108 | 99 | 94 | #6C635E | Dorado |
| 82 | PAP 835 | 41 | 39 | 37 | #292725 | Bokara Grey |
| 83 | PAP 845 | 36 | 32 | 28 | #24201C | Bokara Grey |
| 84 | PAP 932 | 54 | 57 | 65 | #363941 | Vulcan |
| 85 | PAP 1025 | 100 | 89 | 78 | #64594E | Domino |
| 86 | PAP 1062 | 98 | 83 | 66 | #625342 | Judge Grey |
| 87 | PAP 1090 | 94 | 90 | 91 | #5E5A5B | Don Juan |
| 88 | PAP 1129 | 52 | 51 | 55 | #343337 | Haiti |
| 89 | PAP 1142 | 158 | 135 | 108 | #9E876C | Sandal |
| 90 | PAP 1168 | 100 | 95 | 95 | #645F5F | Zambezi |
| 91 | PAP 1185 | 121 | 120 | 123 | #79787B | Topaz |
| 92 | PAP 1224 | 120 | 122 | 118 | #787A76 | Gunsmoke |
| 93 | PAP 180 A | 141 | 116 | 72 | #8D7448 | Shadow |
| 94 | PAP 180 B | 140 | 114 | 69 | #8C7245 | Shadow |
| 95 | PAP 229 A | 34 | 30 | 31 | #221E1F | Aubergine |

**(a)**

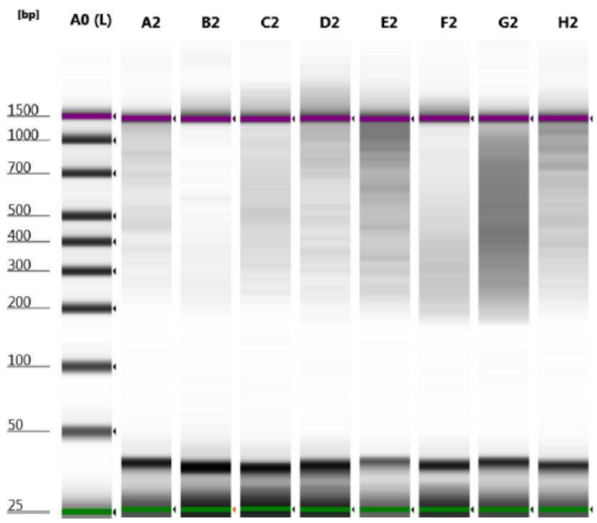


| Gel Position | Enzyme Combination |
|---|---|
| **A2** | PstI/MspI |
| **B2** | PstI/MseI |
| **C2** | PstI/NlaIII |
| **D2** | PstI/HpyCH4IV |
| **E2** | EcoRI/MspI |
| **F2** | EcoRI/MseI |
| **G2** | EcoRI/NlaIII |
| **H2** | EcoRI/HpyCH4IV |

**(b)** ***Pst*I/*Msp*I**

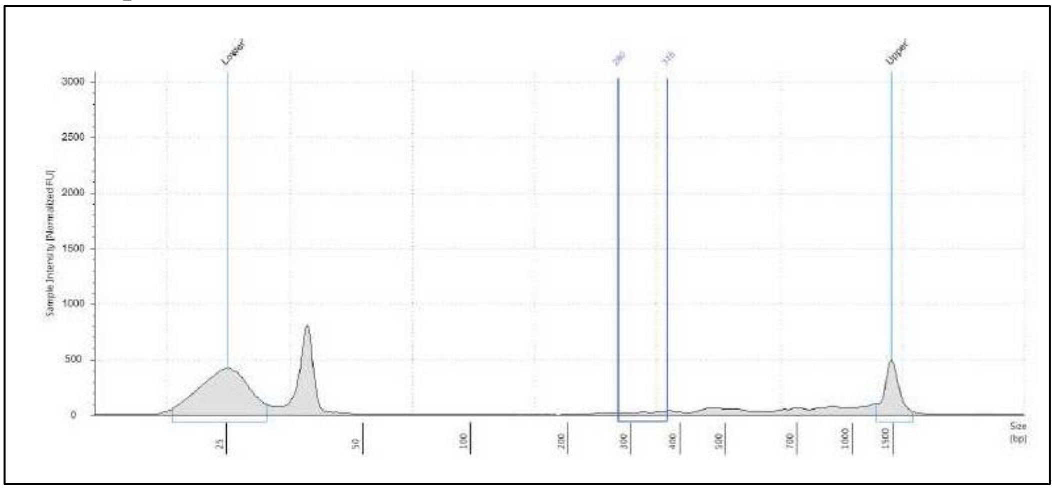


**(c)** ***Pst*I/*Mse*I**

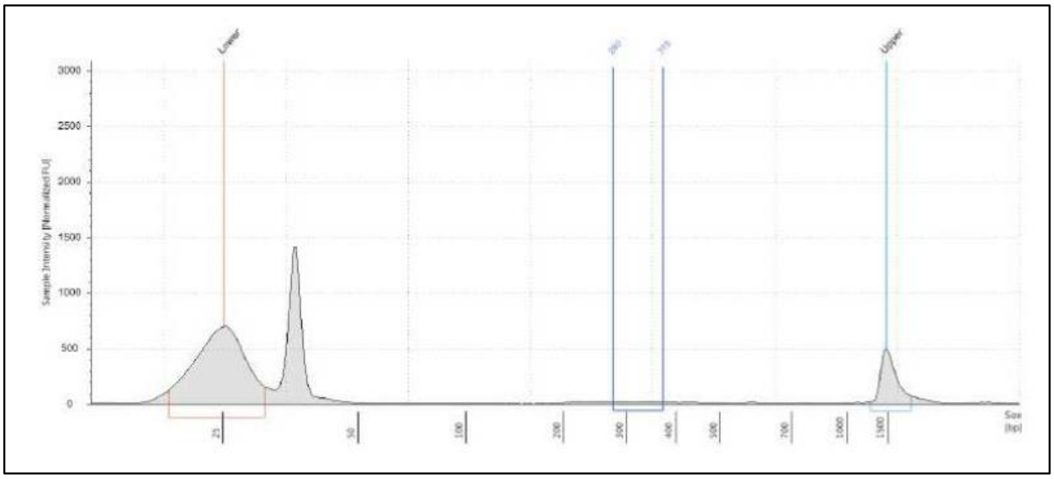


(Fig. **S1** continued)

**(d)** ***Pst*I/*Nla*III**

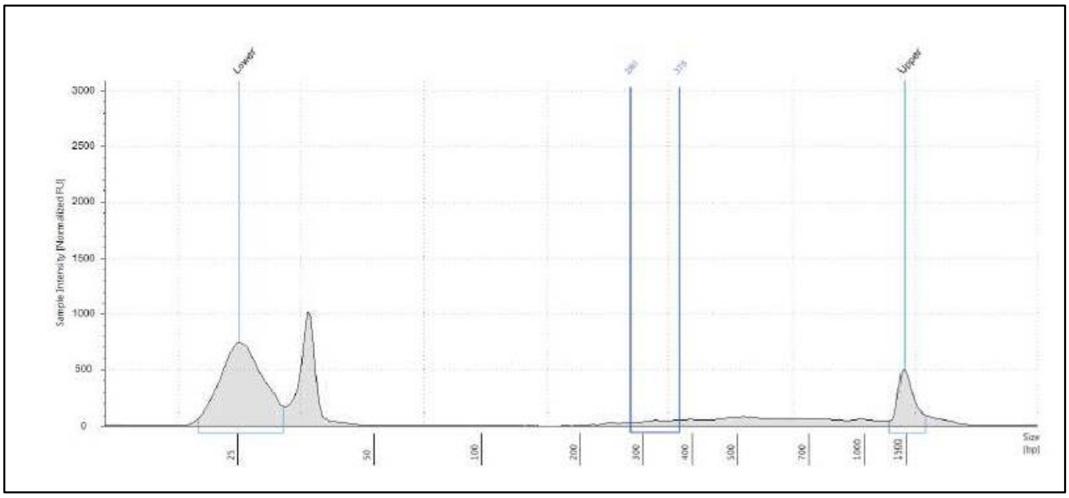


**(e)** ***Pst*I/*HpyCh4*IV**

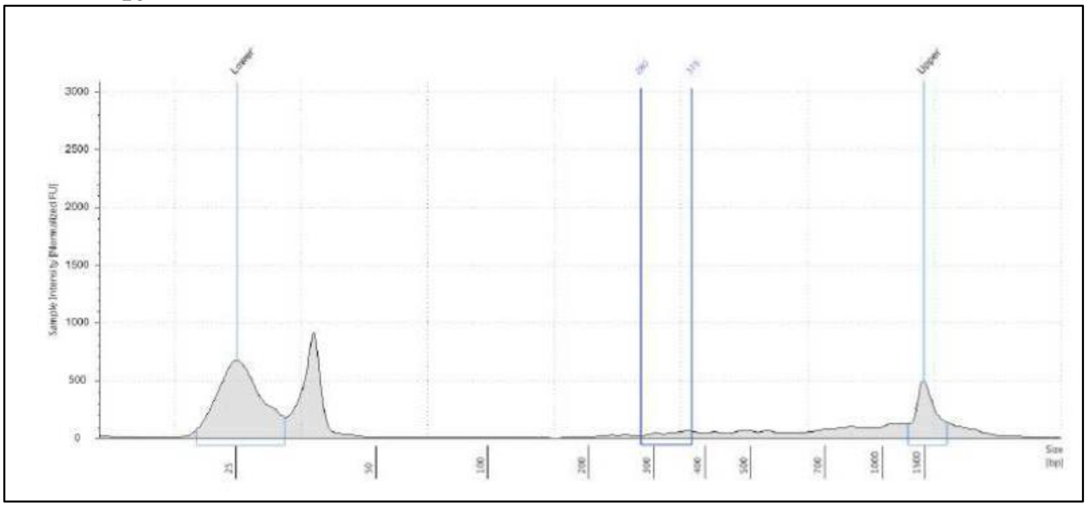


**(f)** ***Eco*RI/*Msp*I**

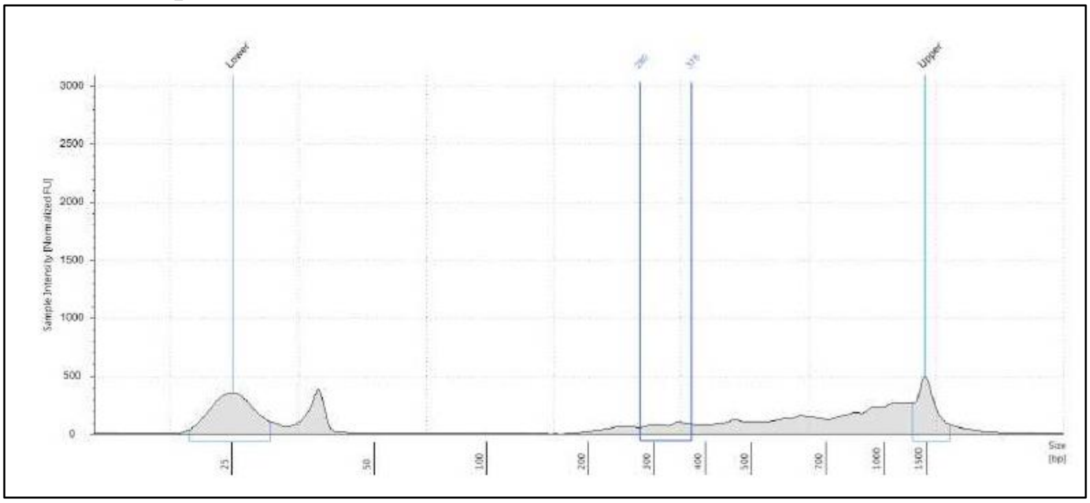


(Fig. **S1** continued)

**(g)** ***Eco*RI/*Mse*I**

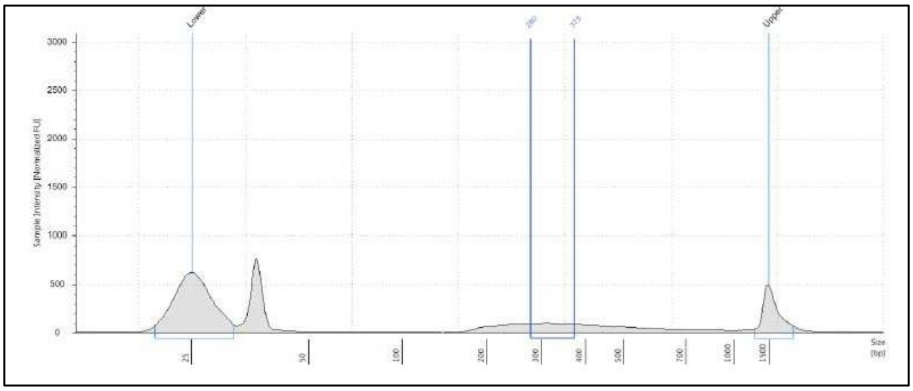


**(h)** ***Eco*RI/*Nla*III**

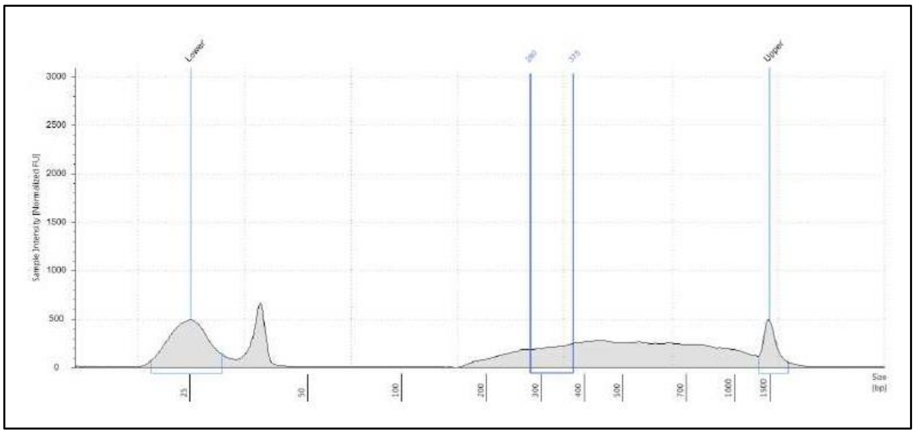


**(i)** *Eco*RI/*HpyCh4*IV

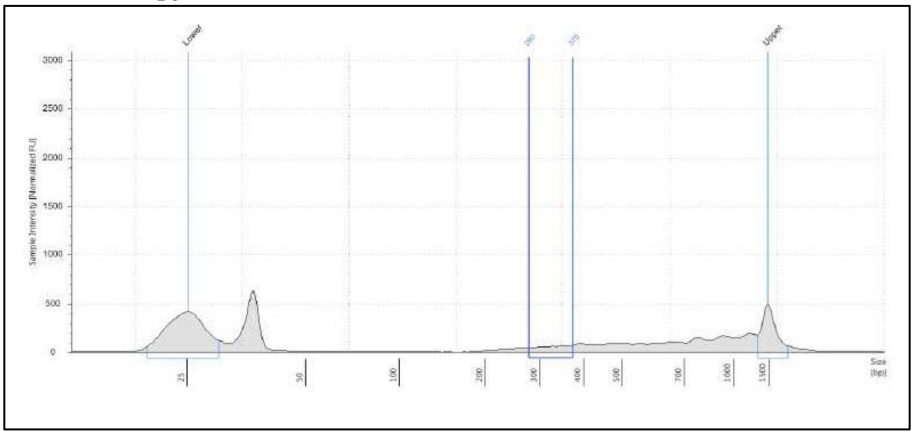


**Fig. S1** Selecting the optimum restriction enzymes combination for GBS in *Papaver* species. (**a**) A gel view of the eighth double digested libraries from a pool of 3 representative samples used for selecting the optimal enzyme combination for GBS in *Papaver*. The eight enzyme combinations used are indicated to the right. (**b-i**) Electropherogram views of the eight double digested libraries from a pool of 3 representative samples used for selecting the optimal enzyme combination for GBS in *Papaver*. An even profile within the size selection window (280bp-375bp) means no or low repeat regions, which usually appear as a peak in the electropherogram view. The enzyme combination used for each electropherogram is show at the top.

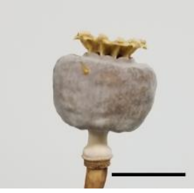
PAP 400
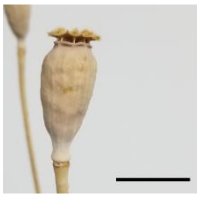
PAP 796
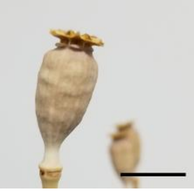
PAP 229 A
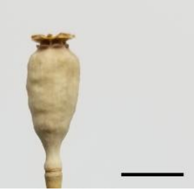
PAP 255
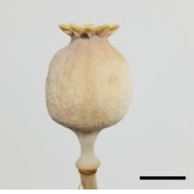
PAP 354
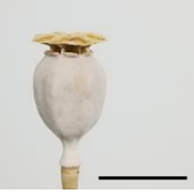
PAP 696
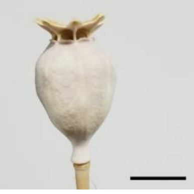
PAP 739
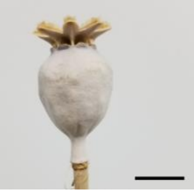
PAP 749
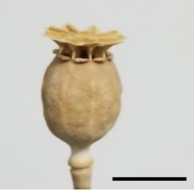
PAP 333
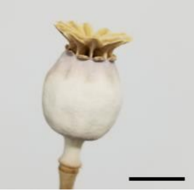
PAP 835
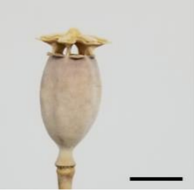
PAP 478
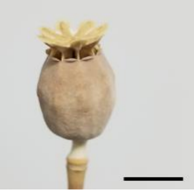
PAP 452
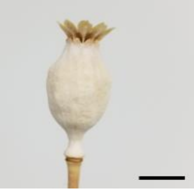
PAP 814
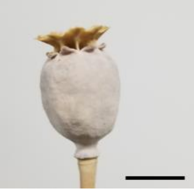
PAP 518
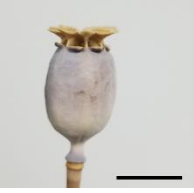
PAP 715
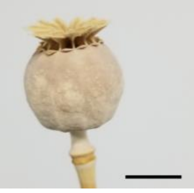
PAP 337
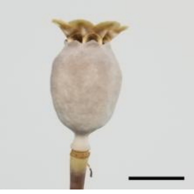
PAP 336
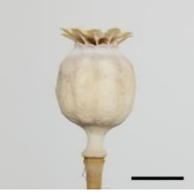
PAP 795
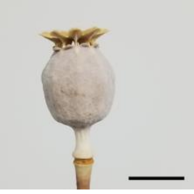
PAP 496
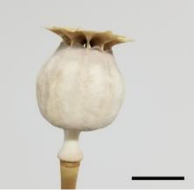
PAP 777
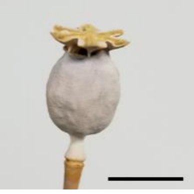
PAP 823
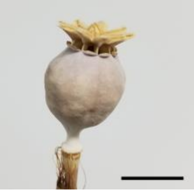
PAP 469
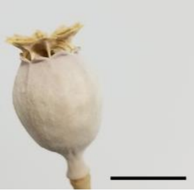
PAP 816
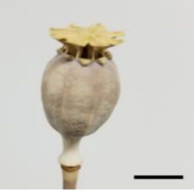
PAP 815
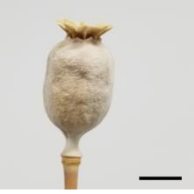
PAP 845
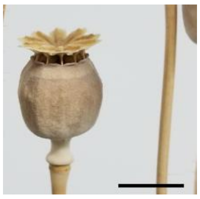
PAP 683
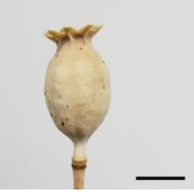
PAP 1142
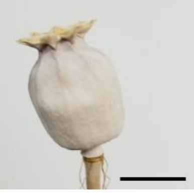
PAP 8
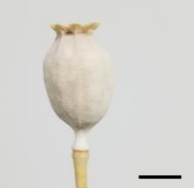
PAP 764
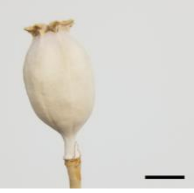
PAP 765
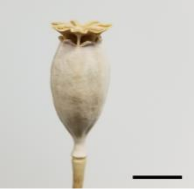
PAP 767
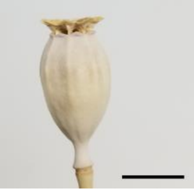
PAP 766

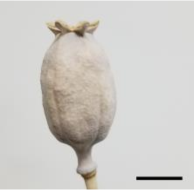
PAP 142
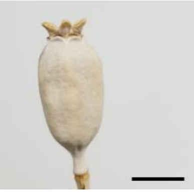
PAP 790
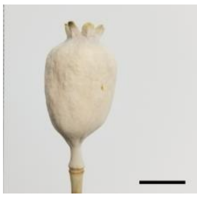
PAP 140
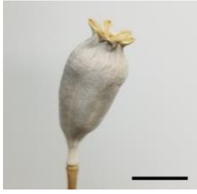
PAP 59
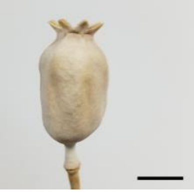
PAP 327
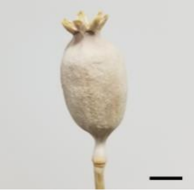
PAP 149
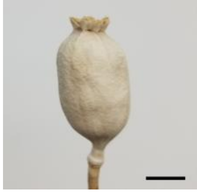
PAP 151
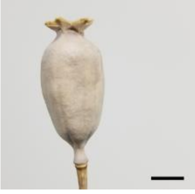
PAP 150
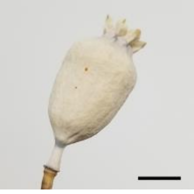
PAP 200
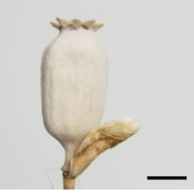
PAP 761
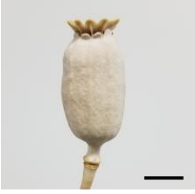
PAP 630
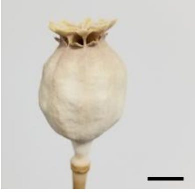
PAP 466
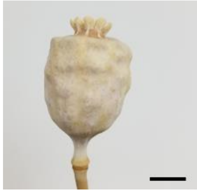
PAP 167
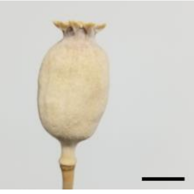
PAP 1090
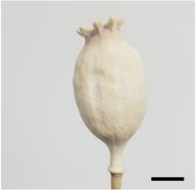
PAP 281
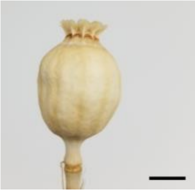
PAP 28
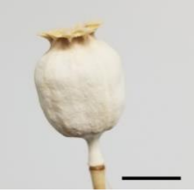
PAP 1224
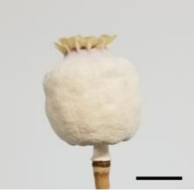
PAP 1185
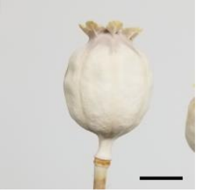
PAP 160
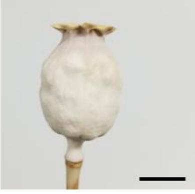
PAP 12
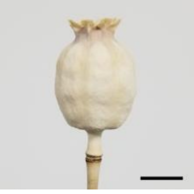
PAP 1129
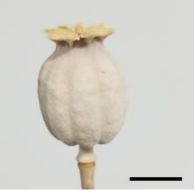
PAP 107
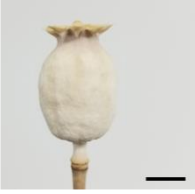
PAP 692
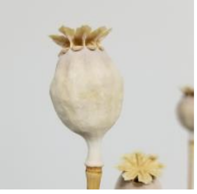
PAP 483
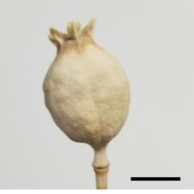
PAP 733
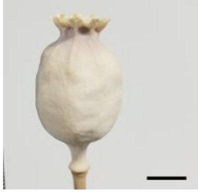
PAP 599
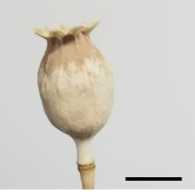
PAP 56
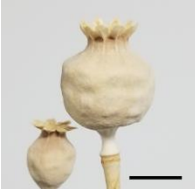
PAP 335
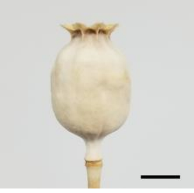
PAP 612
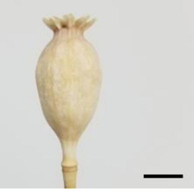
PAP 784
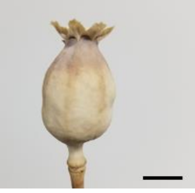
PAP 201
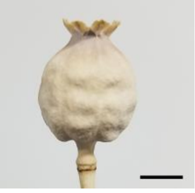
PAP 393

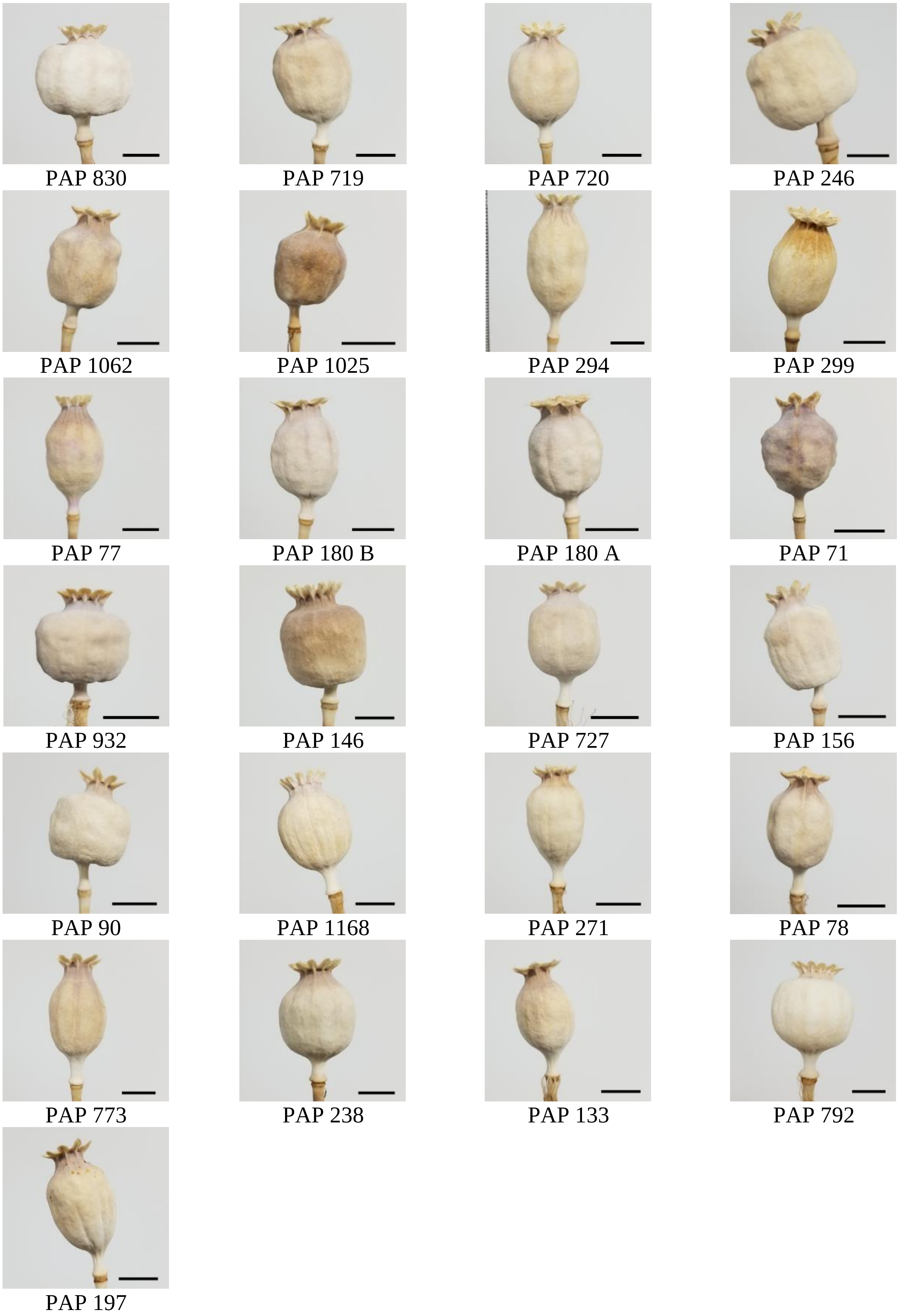


**Fig. S2** Capsule morphology of the *Papaver* accessions included in this study. Bar = 1 cm. Images were taken using a SM-G950U1 digital camera under normal light indoors.

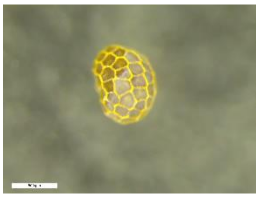
PAP 400

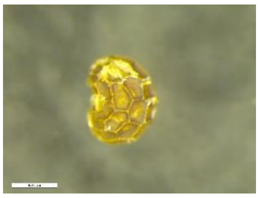
PAP 796

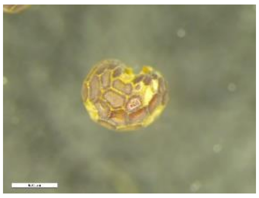
PAP 229 A

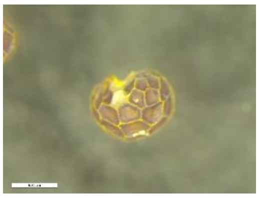
PAP 255

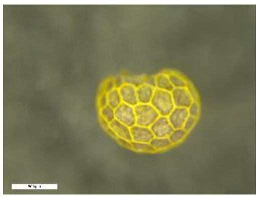
PAP 354

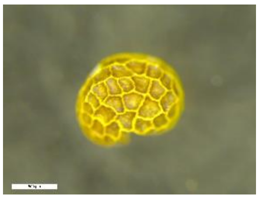
PAP 696

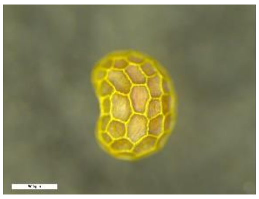
PAP 739

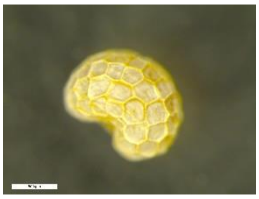
PAP 749

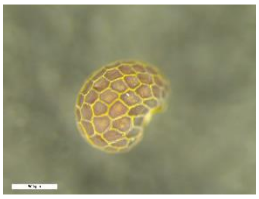
PAP 333

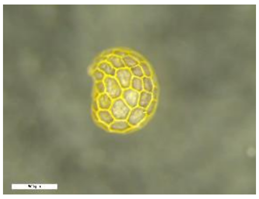
PAP 835

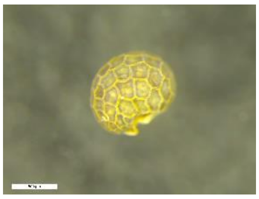
PAP 478

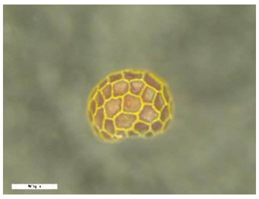
PAP 452

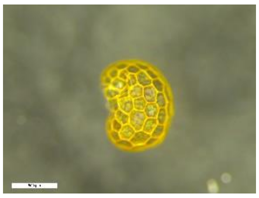
PAP 814

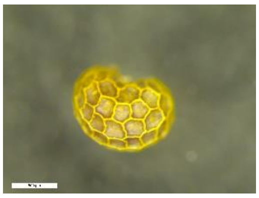
PAP 518

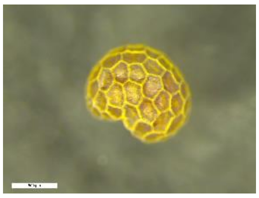
PAP 715

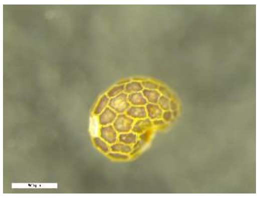
PAP 337

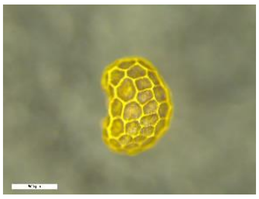
PAP 336

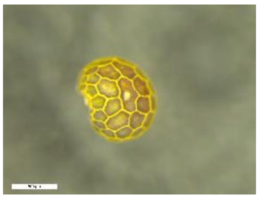
PAP 795

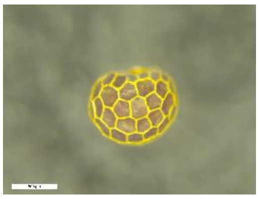
PAP 496

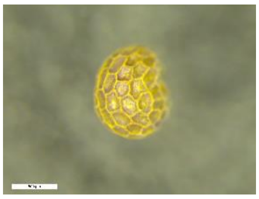
PAP 777

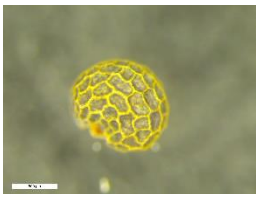
PAP 823

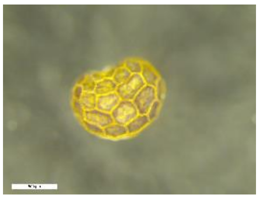
PAP 469

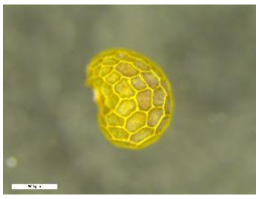
PAP 816

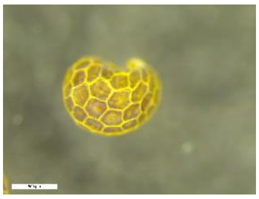
PAP 815

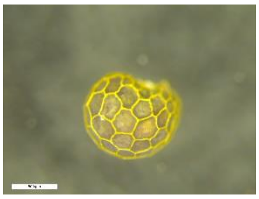
PAP 845

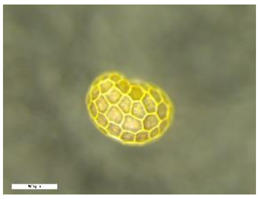
PAP 683

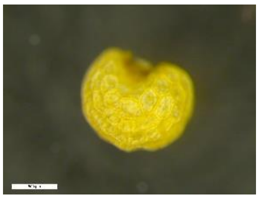
PAP 1142

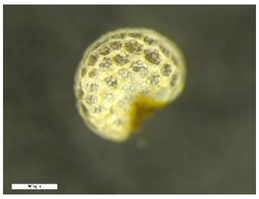
PAP 8

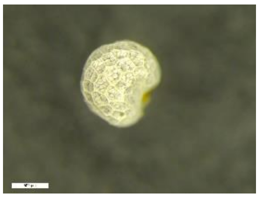
PAP 764

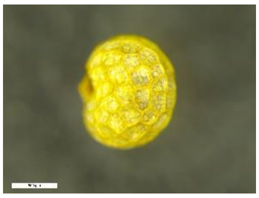
PAP 765

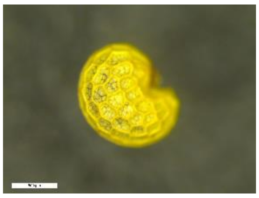
PAP 767

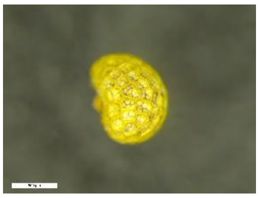
PAP 766

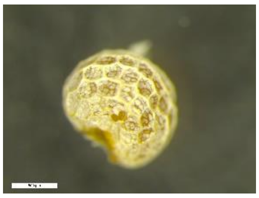
PAP 142
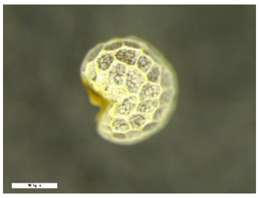
PAP 790
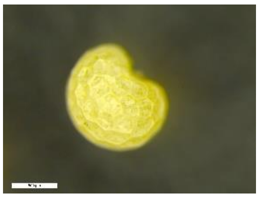
PAP 140
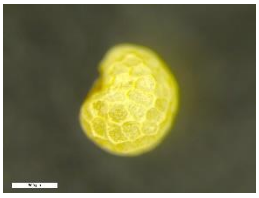
PAP 59
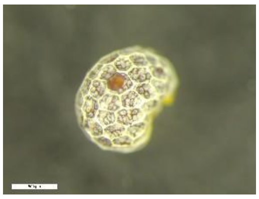
PAP 327
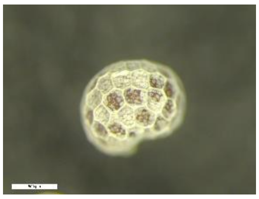
PAP 149
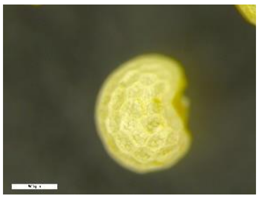
PAP 151
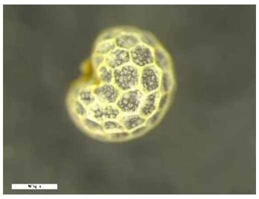
PAP 150
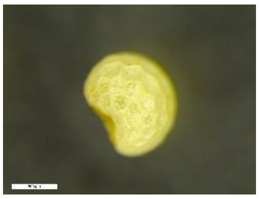
PAP 200
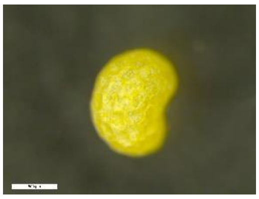
PAP 761
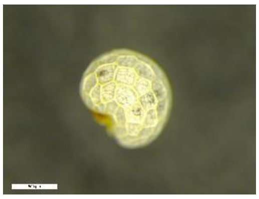
PAP 630
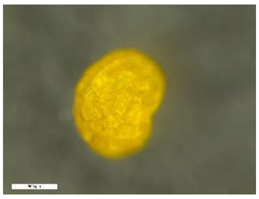
PAP 466
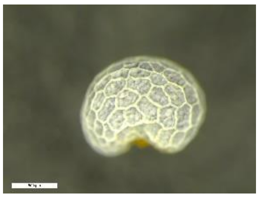
PAP 167
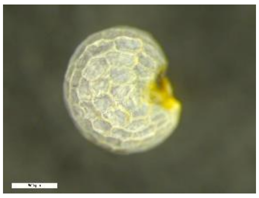
PAP 1090
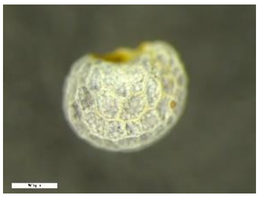
PAP 281
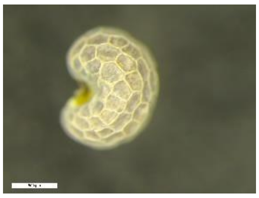
PAP 28
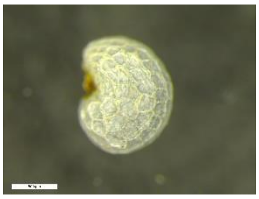
PAP 1224
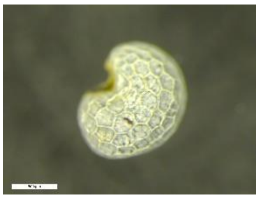
PAP 1185
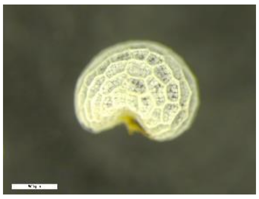
PAP 160
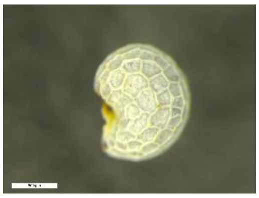
PAP 12
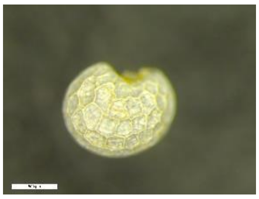
PAP 1129
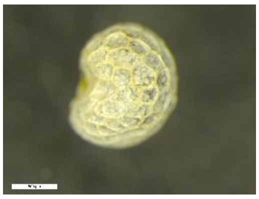
PAP 107
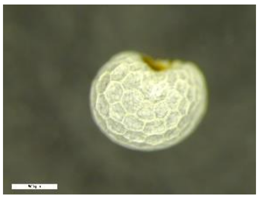
PAP 692
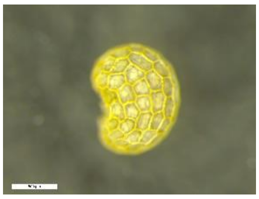
PAP 483
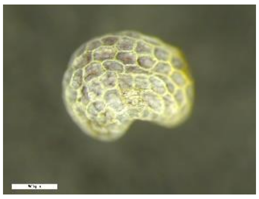
PAP 733
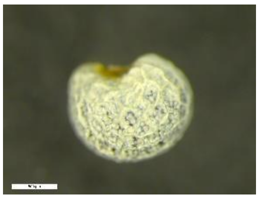
PAP 599
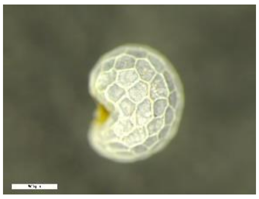
PAP 56
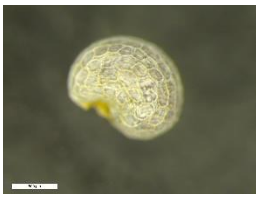
PAP 335
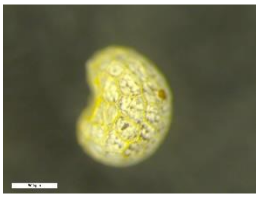
PAP 612
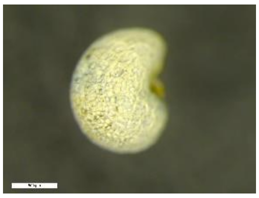
PAP 784
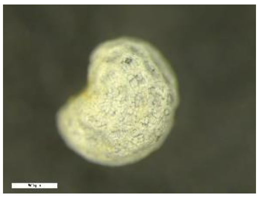
PAP 201
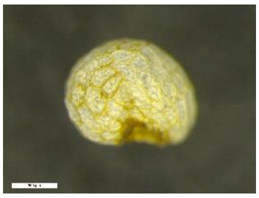
PAP 393

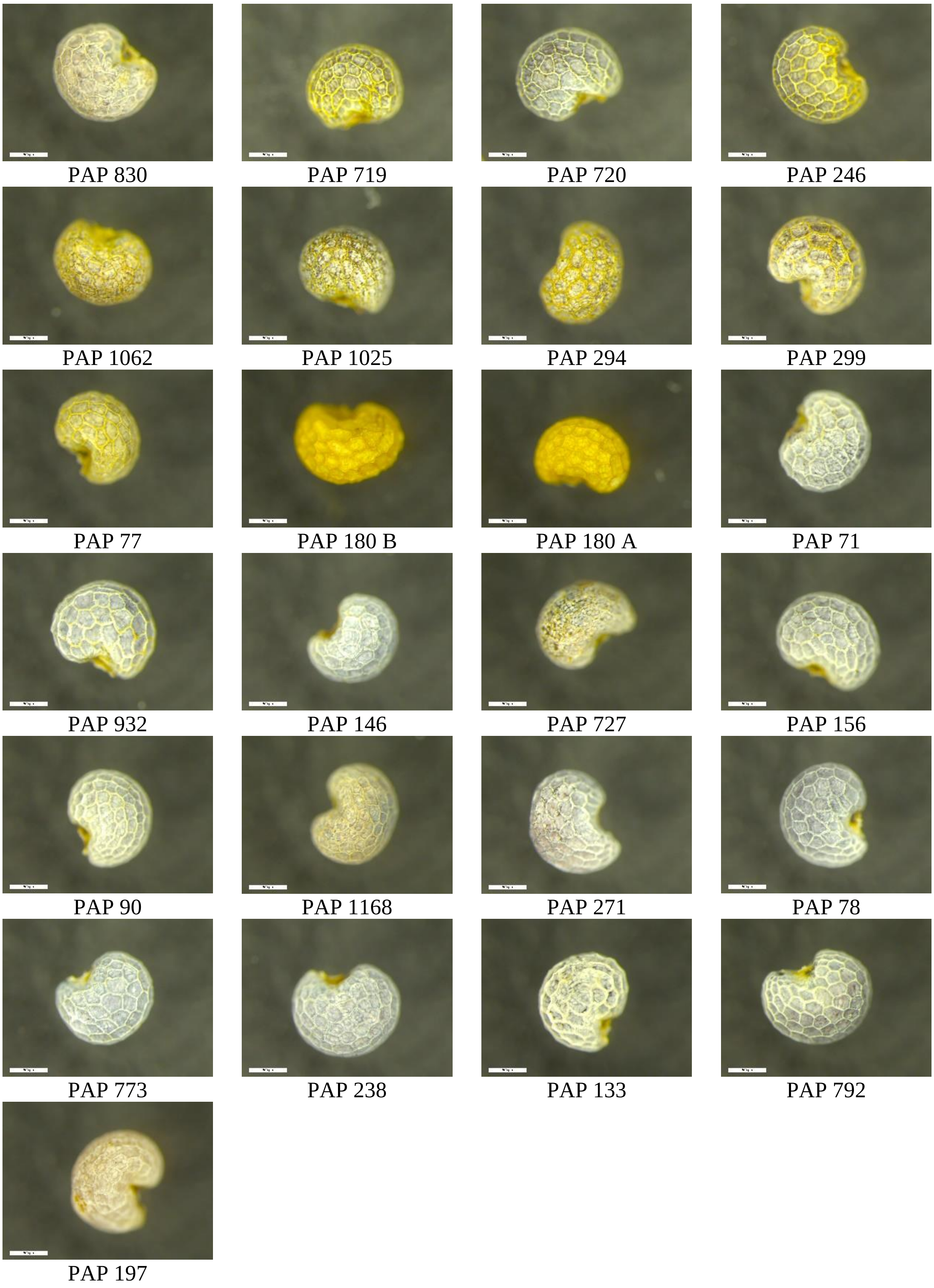


**Fig. S3** Seed morphology of the *Papaver* accessions included in this study. Bar = 500 μm. Images were captured using a Leica M205 FA stereomicroscope with a digital camera Leica DFC 420 and Leica Application Suite Software (LAS 4.0, Leica) with 10% light and 50% zoom in condition.

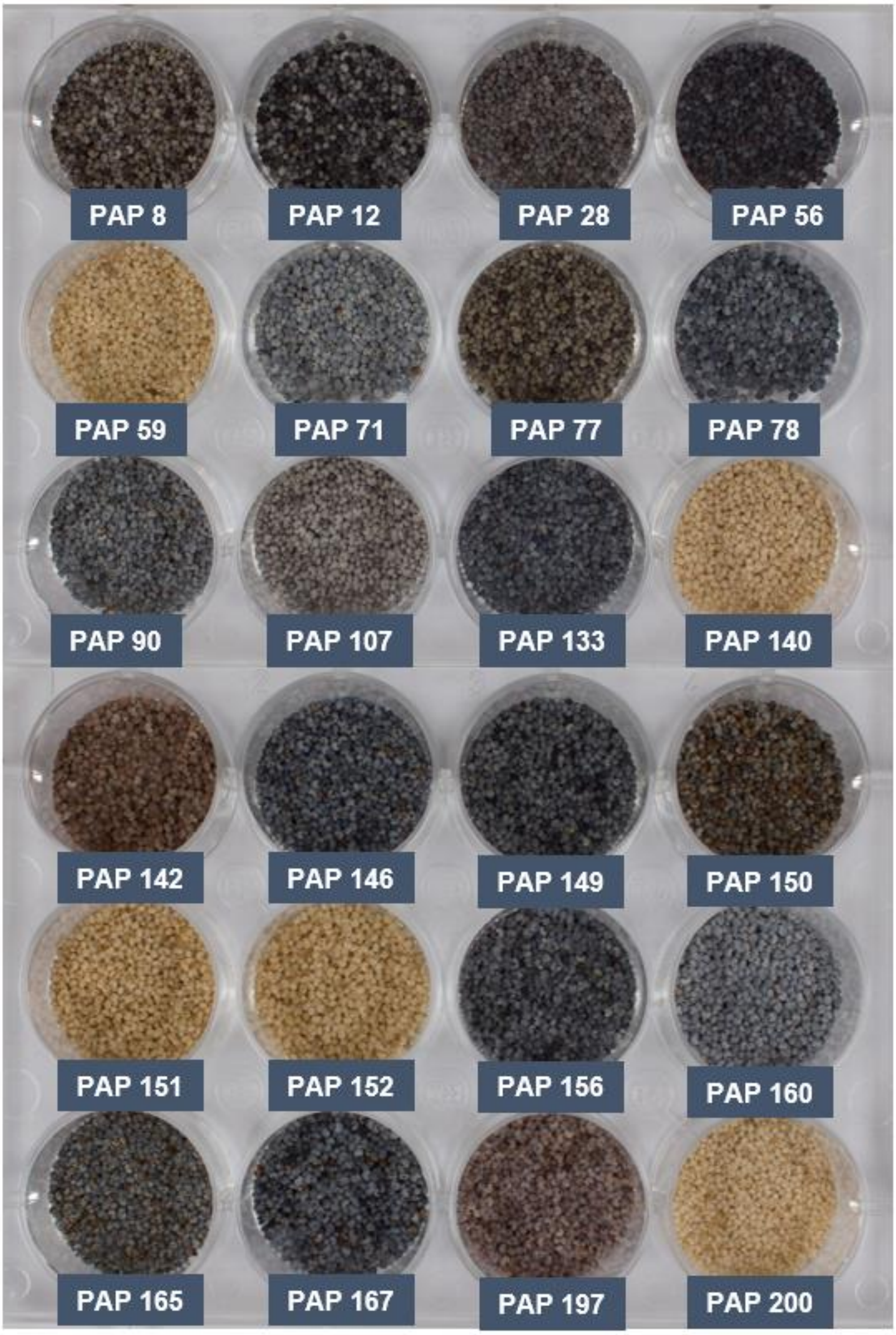


(Fig. **S4** continued)

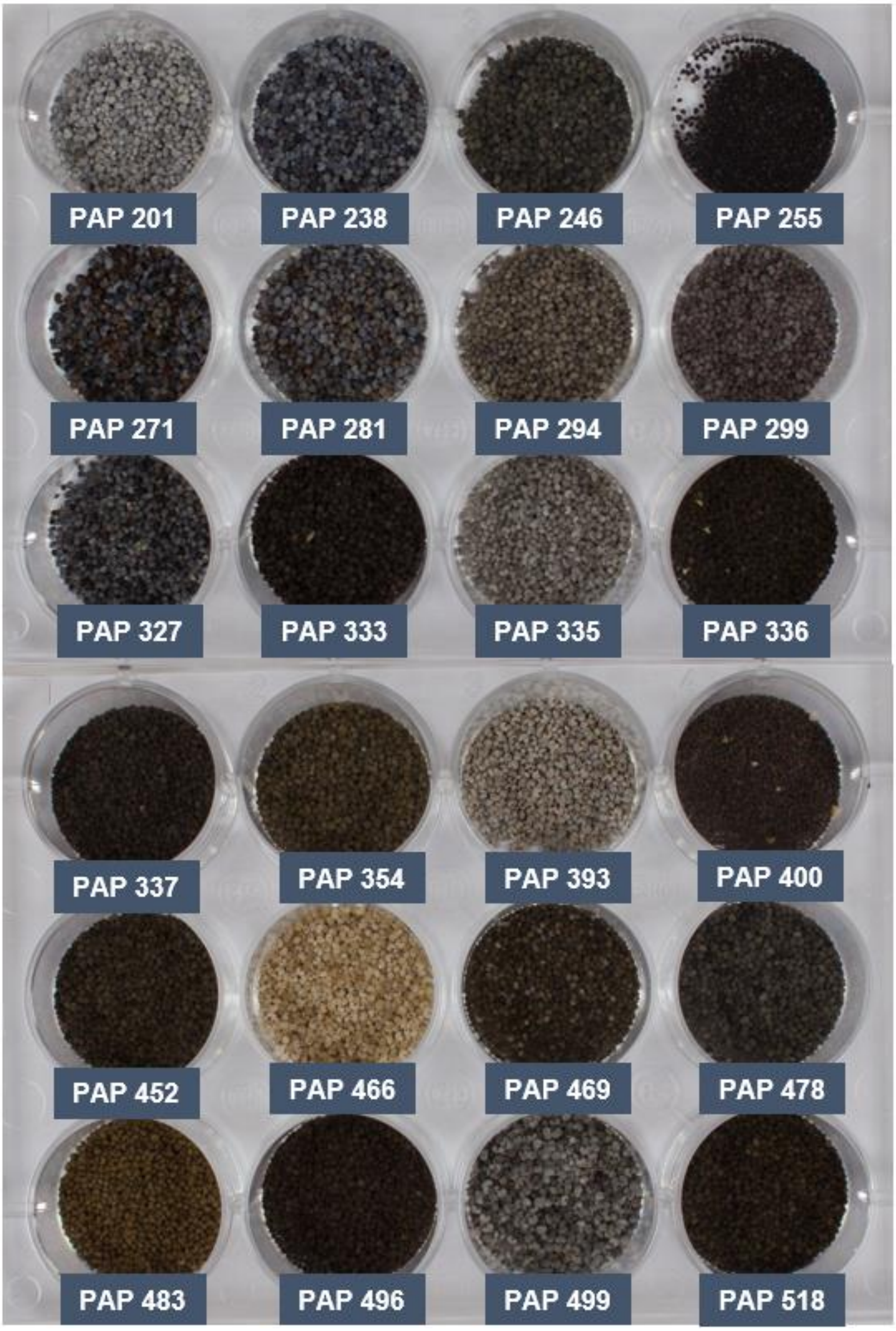


(Fig. **S4** continued)

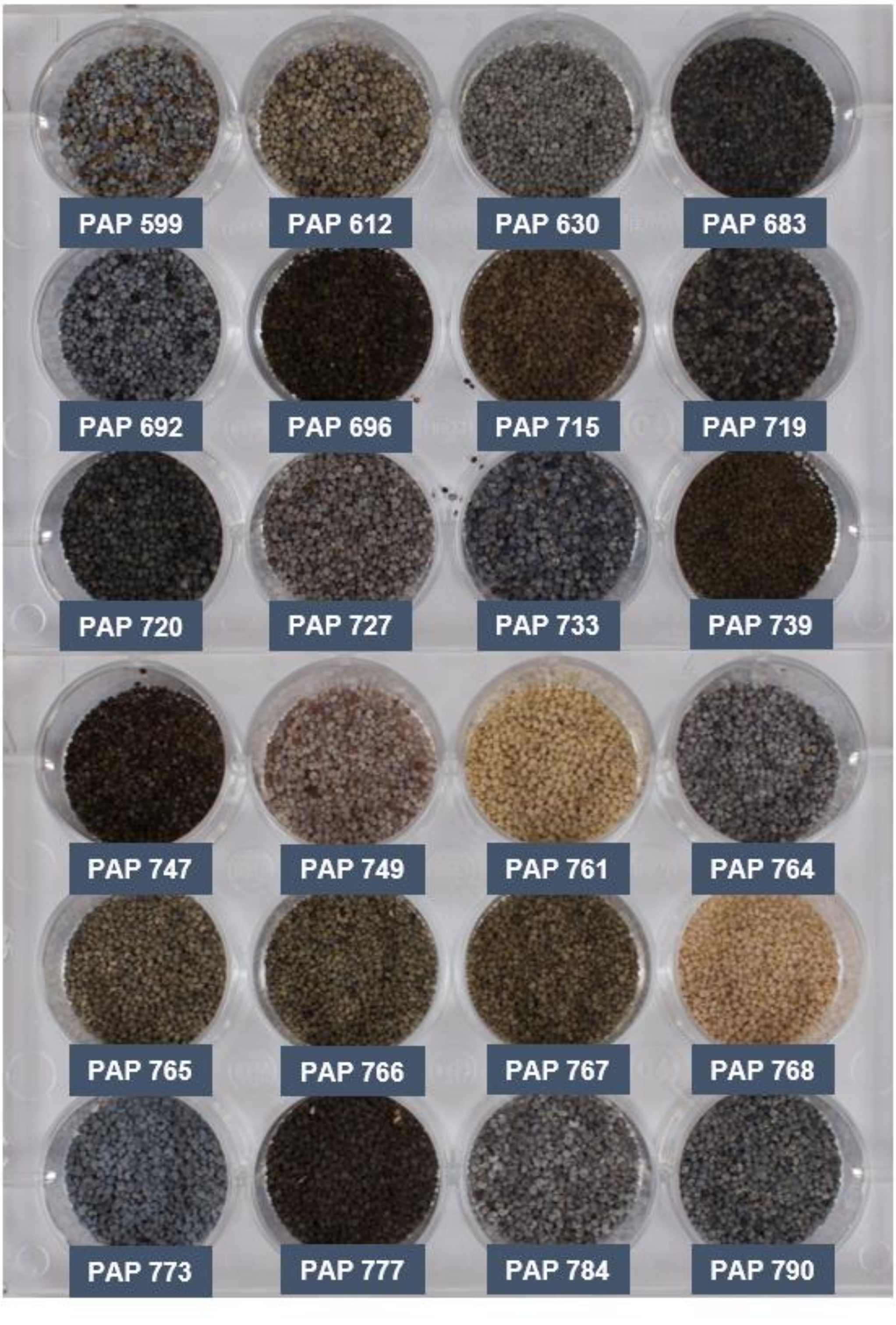


(Fig. **S4** continued)

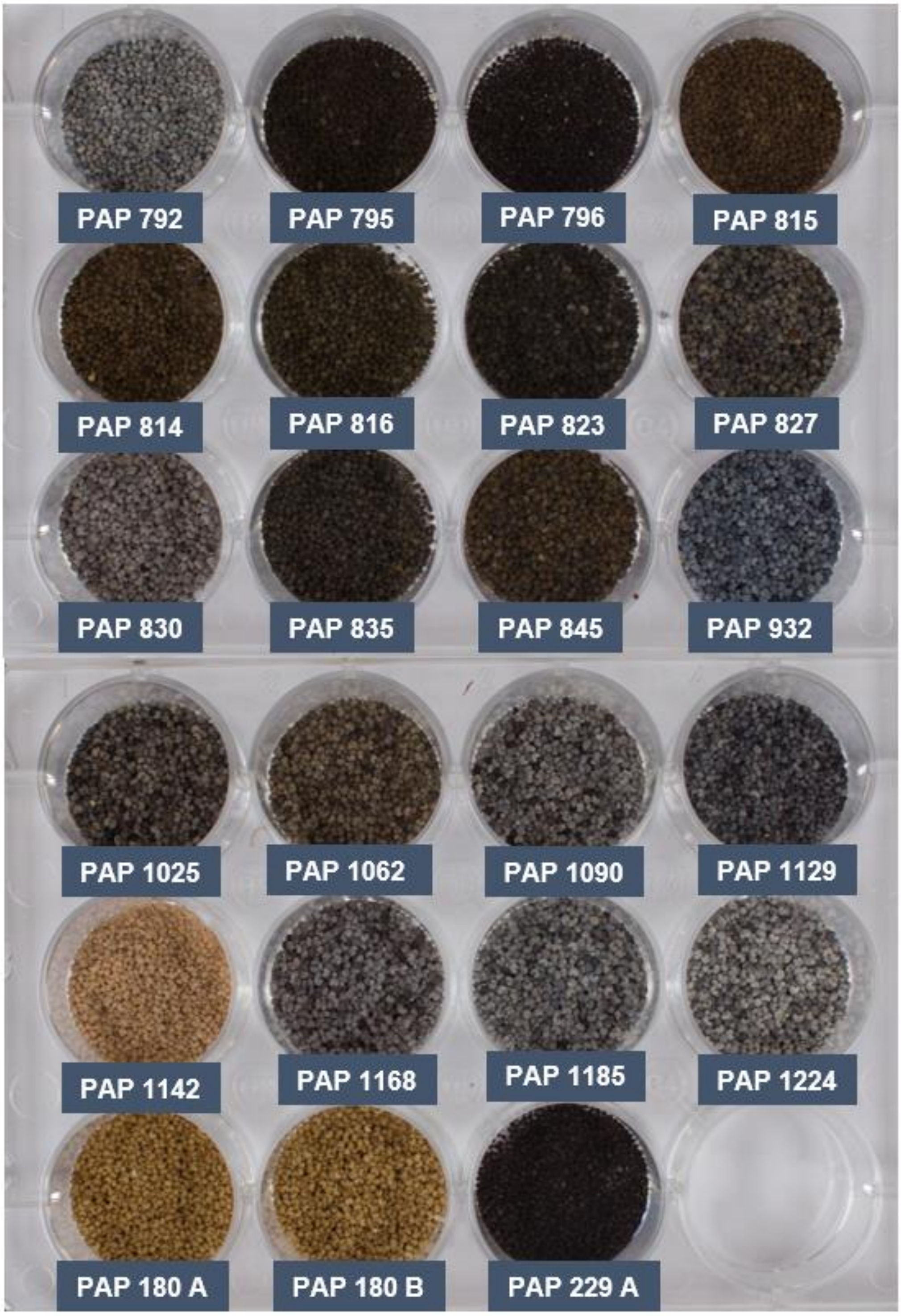


**Fig. S4** Seed colour variability of the *Papaver* accessions included in this study. The RGB values were estimated from raw pictures using ColorChecker (X-rite) in Adobe Lightroom Classic CC software. The RGB values were converred to Hex colour codes, from which the corresponsding approximate colour were determined using HTML CSS Color Picker (see **Table S6**).

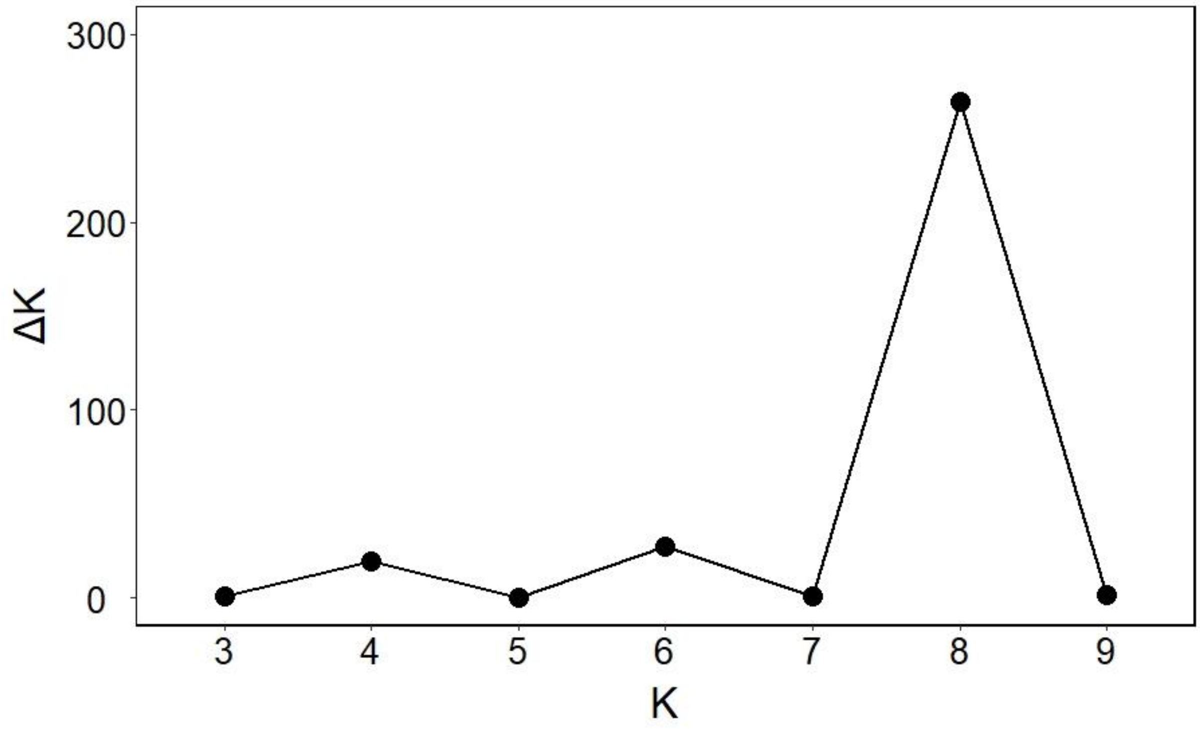


**Fig. S5** Plot of ΔK values with K from 3 to 9 obtained from the STRUCTURE analysis.

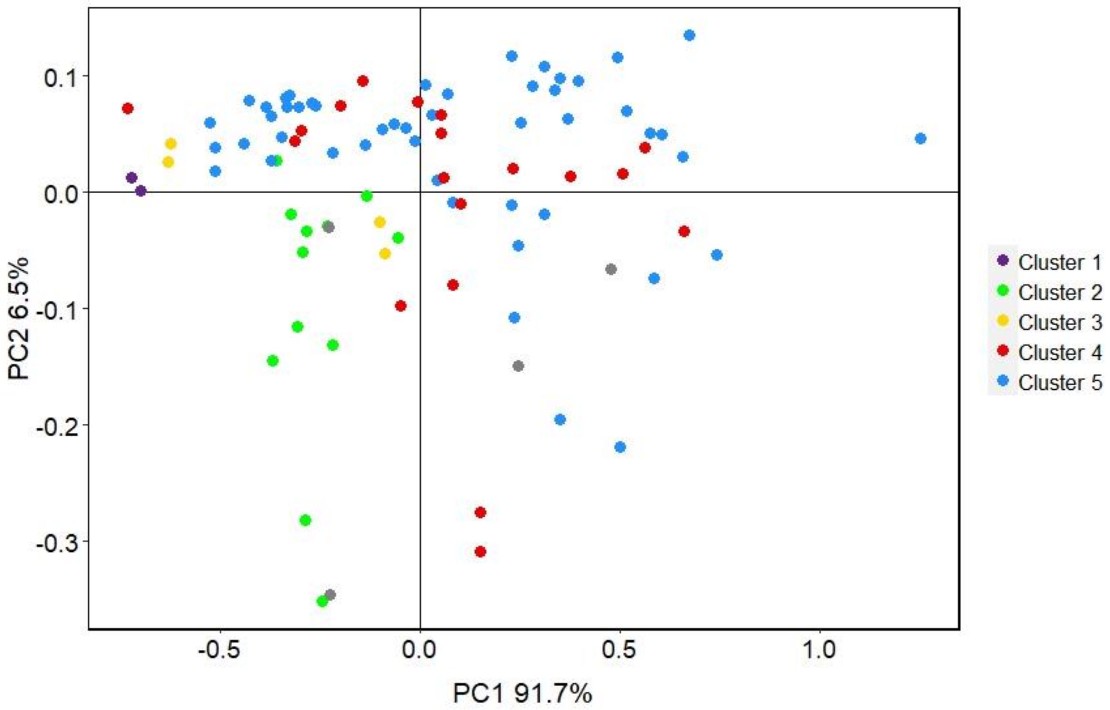


**Fig. S6** A two-dimensional PCA score plot showing the diversity in alkaloid content. Axes are labelled with the percentage of variation explained by the two PCs. Colours indicate the five subpopulations identified by DAPC (**Fig. 5**). The accessions for which no GBS data was available are indicated with grey dots.
